# Cerebellar plasticity and associative memories are controlled by perineuronal nets

**DOI:** 10.1101/736694

**Authors:** Daniela Carulli, Robin Broersen, Fred de Winter, Elizabeth M. Muir, Maja Mešković, Matthijs de Waal, Sharon de Vries, Henk-Jan Boele, Cathrin B. Canto, Chris I. De Zeeuw, Joost Verhaagen

## Abstract

Perineuronal nets (PNNs) are assemblies of extracellular matrix molecules, which surround the cell body and dendrites of many types of neuron and regulate neural plasticity. PNNs are prominently expressed around neurons of the deep cerebellar nuclei (DCN) but their role in adult cerebellar plasticity and behavior is far from clear. Here we show that PNNs in the DCN are diminished during eyeblink conditioning (EBC), a form of associative motor learning that depends on DCN plasticity. When memories are fully acquired, PNNs are restored. Enzymatic digestion of PNNs in the DCN improves EBC learning, but intact PNNs are necessary for memory retention. At the structural level, PNN removal induces significant synaptic rearrangements *in vivo*, resulting in increased inhibition of DCN baseline activity in awake behaving mice. Together, these results demonstrate that PNNs are critical players in the regulation of cerebellar circuitry and function.

## Introduction

Perineuronal nets (PNNs) are lattice-like aggregates of extracellular matrix molecules surrounding the cell body and dendrites of various types of neurons in the central nervous system (CNS). Synaptic contacts impinging on PNN-bearing neurons are embedded in these structures. Therefore, PNNs are strategically positioned to influence the development and stabilization of synaptic connections. PNNs emerge during late postnatal development, in coincidence with the closure of critical periods for experience-dependent plasticity (Takesian and Hensch, 2013; Fawcett et al., 2019). Accumulating evidence suggests that PNNs inhibit different forms of CNS plasticity in adult life, both in physiological and pathological conditions. For instance, enzymatic digestion of PNNs with chondroitinase ABC (ch’ase) or manipulation of PNN components enhance cortical plasticity (Pizzorusso et al., 2002; Carulli et al., 2010; Bernard and Prochiantz, 2016; Lee et al., 2017; Rowlands et al., 2018; Boggio et al., 2019), cognitive flexibility (Happel et al., 2014), AMPA receptor-dependent synaptic plasticity (Frischknecht et al., 2009), and axonal sprouting after injury (Galtrey et al., 2007; Massey et al., 2006; Carulli et al., 2010; Bowes et al., 2012; Soleman et al., 2012; Gherardini et al., 2015). PNN disruption also increases the formation of recognition memory (Romberg et al., 2013, Yang et al., 2015), whereas it impairs the consolidation of other types of memory (Hylin et al., 2013; Xue et al., 2014; Slaker et al., 2015; Banerije et al., 2017; Carbo-Gas et al., 2017; Blacktop et al., 2017; Thompson et al., 2018; Blacktop and Sorg, 2019). Ch’ase digests chondroitin sulfate proteoglycans (CSPGs), one of the major components of the PNN, which consist of a core protein and a variable number of glycosaminoglycan side chains. The sulfation pattern of glycosaminoglycans is critical for PNN function, as it encodes specific information for the binding of plasticity regulators (Miyata et al., 2012; Beurdeley et al., 2012; Vo et al., 2013; Dick et al., 2013; Hou et al., 2017).

The cerebellum shows abundant PNNs around neurons in the deep cerebellar nuclei (DCN) (Carulli et al., 2006). The cerebellum plays a pivotal role in motor, emotional and cognitive associative learning (Timmann et al., 2010; Gao et al., 2012; Rochefort et al., 2013). Delay eyeblink conditioning (EBC) is an associative learning paradigm, which consists of an eyelid closure in response to a previously neutral stimulus (such as a light; conditioned stimulus - CS) after repeated pairing of the CS with an obligatory eyeblink-eliciting stimulus (such as an air puff; unconditioned stimulus - US). EBC critically depends on DCN function (Freeman and Steinmetz, 2011). CS and US inputs reach the DCN via excitatory collaterals of mossy fibers originating in the basilar pontine nuclei and of climbing fibers originating in the inferior olive, respectively (De Zeeuw and Yeo, 2005). While mossy fibers can excite Purkinje cells in the cerebellar cortex through their innervation of the granule cells that give rise to the parallel fibers, the climbing fibers excite the Purkinje cells directly. Purkinje cells in turn inhibit the DCN neurons (Ito et al., 1972), allowing an integration of the excitatory collaterals with inhibitory inputs (Pugh and Raman, 2009). At the end of EBC the activity of DCN neurons is enhanced during expression of the conditioned response (CR) (Ten Brinke et al., 2017). In spite of the presence of PNNs, multiple forms of synaptic and structural plasticity take place in the DCN during EBC (Kleim et al., 2002; Freeman, 2015; Boele et al., 2013). Indeed, following EBC the amplitude of the CR can be correlated with the density of mossy fiber terminals derived from the pontine nuclei (Boele et al., 2013), raising the possibility that the PNNs in the DCN need to be reduced so as to allow sprouting of the mossy fiber collaterals.

In the current study, we sought to unravel the role of PNNs in the control of cerebellar plasticity at the circuit and behavioral level. We investigated whether expression of PNN-CSPGs in the DCN is altered during acquisition and consolidation of EBC in adult mice. Moreover, we overexpressed ch’ase in the DCN via a lentiviral vector and assessed whether PNN-CSPG digestion impacts the performance of mice both during and after formation of EBC memories. Finally, to establish a link between behavioural changes and circuit plasticity, we examined the effects of PNN digestion on remodeling of inhibitory and excitatory synaptic terminals and on the baseline electrophysiological properties of DCN neurons *in vivo*. Our data provide the first evidence for a dynamic modulation of PNNs in response to EBC and for a tight control of the balance between excitatory and inhibitory inputs to DCN neurons by PNNs, thereby regulating acquisition and retention of EBC memory.

## Results

### PNNs are dynamically regulated during eyeblink conditioning

To investigate the role of PNNs in cerebellum-dependent associative learning, we first examined the expression of PNNs enwrapping cerebellar nuclei neurons over the course of EBC. We hypothesized that PNNs are modulated in response to EBC. We examined PNNs during two phases, namely during learning (day 5), and when animals had reached stable levels of performance (day 10; Heiney et al., 2014; Rasmussen et al., 2018). PNN expression was evaluated by quantifying the staining intensity of *Wisteria floribunda* agglutinin (WFA) around individual neurons in eyeblink-encoding regions of the DCN, *i.e*. the dorsolateral hump (DLH) and the adjacent lateral part of the anterior interpositus (IntA) (Yeo et al., 1985; Clark et al., 1992; Krupa and Thompson, 1997; Freeman et al., 2005; Heiney et al., 2014; ten Brinke et al., 2017), with the lateral nucleus as control area (Fig. 1A, B). We found a significant increase in CRs (from 1% on day 1 to 60% on day 5) over the course of 5 days-training in the conditioned group, whereas, as expected, pseudoconditioned animals did not show learning [day: F(1.971, 15.767) = 13.334, P<0.001; group: F(1, 8) = 17.332, P=0.003; interaction: F(1.971, 15.767) = 13.337, P<0.001; Greenhouse-Geisser corrected repeated-measures (RM) ANOVA; Fig. 1D]. Conditioned mice also displayed an increase in the fraction eyelid closure (FEC) at US onset, in contrast to pseudoconditioned mice [day: F(1.746, 13.965) = 11.229, P=0.002; group: F(1, 8) = 14.523, P=0.005; interaction: F(1.746, 13.965) = 10.157, P=0.002; Greenhouse-Geisser corrected RM ANOVA; Fig. 1E]. Since neurons of the DCN display natural variability in WFA staining intensity, we divided the PNNs in three categories: strong, medium, and weak nets (as in Foscarin et al. 2011). Data from both ipsi- and contralateral cerebellar hemispheres (relative to US) were analyzed, because both sides encode EBC (Ivarsson and Hesslow, 1993) and axonal plasticity has been reported for both sides (Boele et al., 2013). WFA intensity distributions were not different between the right and left side in any of the three subnuclei (DLH, IntA and lateral) in control and in pseudoconditioned mice (DLH: control, X^2^_2_ = 0.79, P=0.67; pseudo, X^2^_2_ = 2.03, P=0.36; IntA: control, X^2^_2_ = 3.82, P=0.15; pseudo, X^2^_2_ = 0.52, P=0.77; lateral nucleus: control, X^2^_2_ = 5.28, P=0.07; pseudo, X^2^_2_ = 1.74, P=0.42). Data from the two sides were therefore pooled together. Moreover, no difference was found between pseudoconditioned and control mice after 5 days of EBC (CTR DLH versus pseudo DLH: X^2^_2_ = 4.24, P = 0.12; CTR IntA versus pseudo IntA: X^2^_2_ = 3.13, P = 0.21; CTR lateral versus pseudo lateral: X^2^_2_ = 2.48, P = 0.29; Fig. 1F, G, I-K, M-O, Q), indicating that unpaired presentation of CS and US does not affect PNN expression in the DCN. Interestingly, both in the IntA and the DLH of 5 days-conditioned mice, the percentage of strong nets significantly decreased on both sides when compared to pseudoconditioned and control mice (DLH: from ~ 85% to ~ 55%; IntA: from ~ 65% to ~ 30%), whereas the percentage of medium and weak nets increased (medium nets, DLH: from ~ 15% to ~ 40%; IntA: from ~ 25% to ~ 50%; weak nets, DLH: from ~1% to ~5%; IntA: from ~ 8% to ~ 16%; CTR DLH versus conditioned DLH: X^2^_2_ = 31.97, P<0.001; CTR IntA versus conditioned IntA: X^2^_2_ = 39.30, P<0.001; pseudo DLH versus conditioned DLH: X^2^_2_ = 40.90, P<0.001; pseudo IntA versus conditioned IntA: X^2^_2_ = 51.95, P<0.001; Fig. 1F-M). In addition, in the DLH of conditioned mice, the decrease in WFA intensity was more pronounced on the right side than on the left side (X^2^_2_ = 6.59, P=0.04; Fig. S1A). No difference between right and left side was detected in the IntA of conditioned mice (X^2^_2_ = 3.75, P=0.15; Fig. S1B). However, the effect of EBC on PNN expression in the lateral nucleus was substantially less remarkable. The percentage of strong nets decreased only by ~5% and was statistically significant only when comparing control and conditioned mice (X^2^_2_ = 7.03, P<0.05; pseudo versus conditioned mice: X^2^_2_ = 3.76, P = 0.15; Fig. 1N-Q).

**Figure 1.**
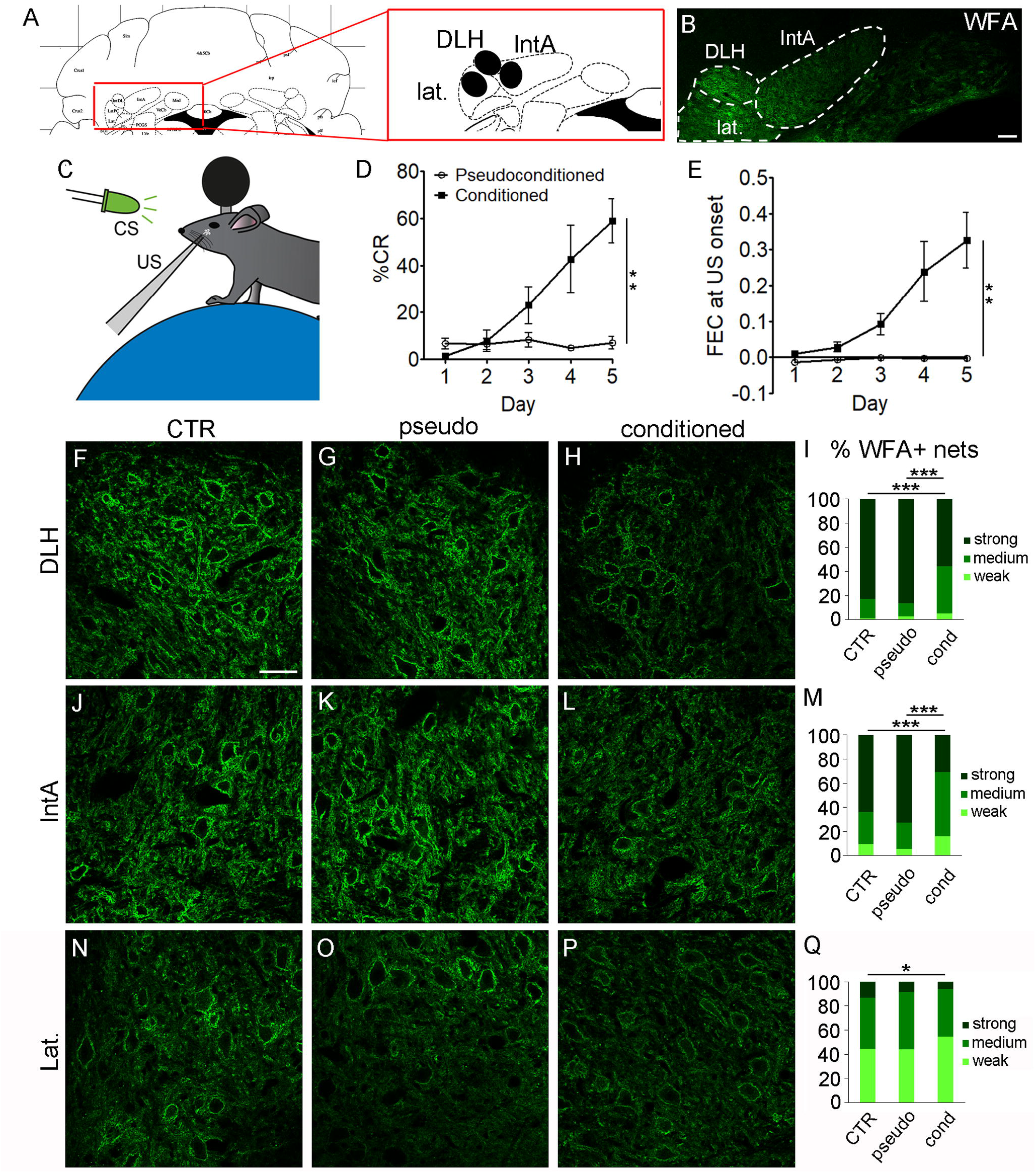
PNNs are reduced during acquisition of EBC. (A) Scheme of a coronal section of the cerebellum [Bregma: −6.12 mm, from Paxinos and Franklin (2001)], showing the location in which the analysis of WFA staining intensity has been performed (black circles). (B) WFA staining in the DCN, where the analyzed nuclei (dorsolateral hump – DLH -, anterior interpositus – IntA -, and lateral nucleus – lat. -) are outlined by dashed lines. (C) Mice were head-fixed on a cylindrical treadmill during EBC training sessions. (D, E) Mice were subjected to EBC (conditioned) or unpaired presentation of CS and US (pseudoconditioned). Learning was observed in the conditioned group as an increase in percentage of trials with CRs (D) and increase in FEC at the time of US onset (E). (F-Q) The intensity of WFA+ nets was analysed in the DLH, the lateral part of the IntA and the lateral nucleus in control (CTR) mice, in mice that were trained for 5 days (conditioned mice), and in pseudoconditioned (pseudo) mice. A significant shift towards weak and medium intensity-nets is apparent in conditioned mice when compared to CTR and pseudo mice in the DLH (F-I) and the IntA (J-M). See also fig. S1. In the lateral nucleus there is only a slight decrease in WFA intensity in conditioned mice when compared to CTR mice (N-Q). There is no difference in the frequency distribution of WFA+ nets between CTR and pseudo animals in all nuclei (I, M, Q). Scale bars: 100 μm in B, 50 μm in F (also applies to G, H, J-L, N-P). * P < 0.05, ** P < 0.01, *** P < 0.001.

To assess whether PNN expression was also altered when animals had completed the acquisition phase, we trained conditioned and pseudoconditioned mice for 10 days. Conditioned mice showed learning as an increase in CRs [from 2% (day 1) to 75% (day 10); day: F(9, 40.452) = 13.941, P<0.001; group: F(1, 9.798) = 104.638, P<0.001; interaction: F(9,40.452) = 11.375, P<0.001, generalized linear mixed model (GLMM); Fig. 2A] and FEC at US onset [day: F(9,40.163) = 9.338, P<0.001; group: F(1,15.301) = 50.729, P<0.001; interaction: F(9,40.163) = 8.514, P<0.001, GLMM; Fig. 2B]. No difference in WFA intensity between right and left sides was detected in the DLH and lateral IntA of conditioned mice (DLH: X^2^_2_ = 1.36, P = 0.51; IntA: X^2^_2_ = 1.47, P = 0.48), thus data from the right and left side were pooled together. In contrast to our findings after 5 days of EBC, after 10 days the WFA intensity distribution in the IntA and the DLH was not different between conditioned and pseudoconditioned mice (DLH: X^2^_2_ = 0.74, P = 0.69; IntA: X^2^_2_ = 0.052, P = 0.97, Fig. 2C-H). These data show that PNNs are transiently reduced during associative motor learning, and are restored when the memory trace has been acquired.

**Figure 2.**
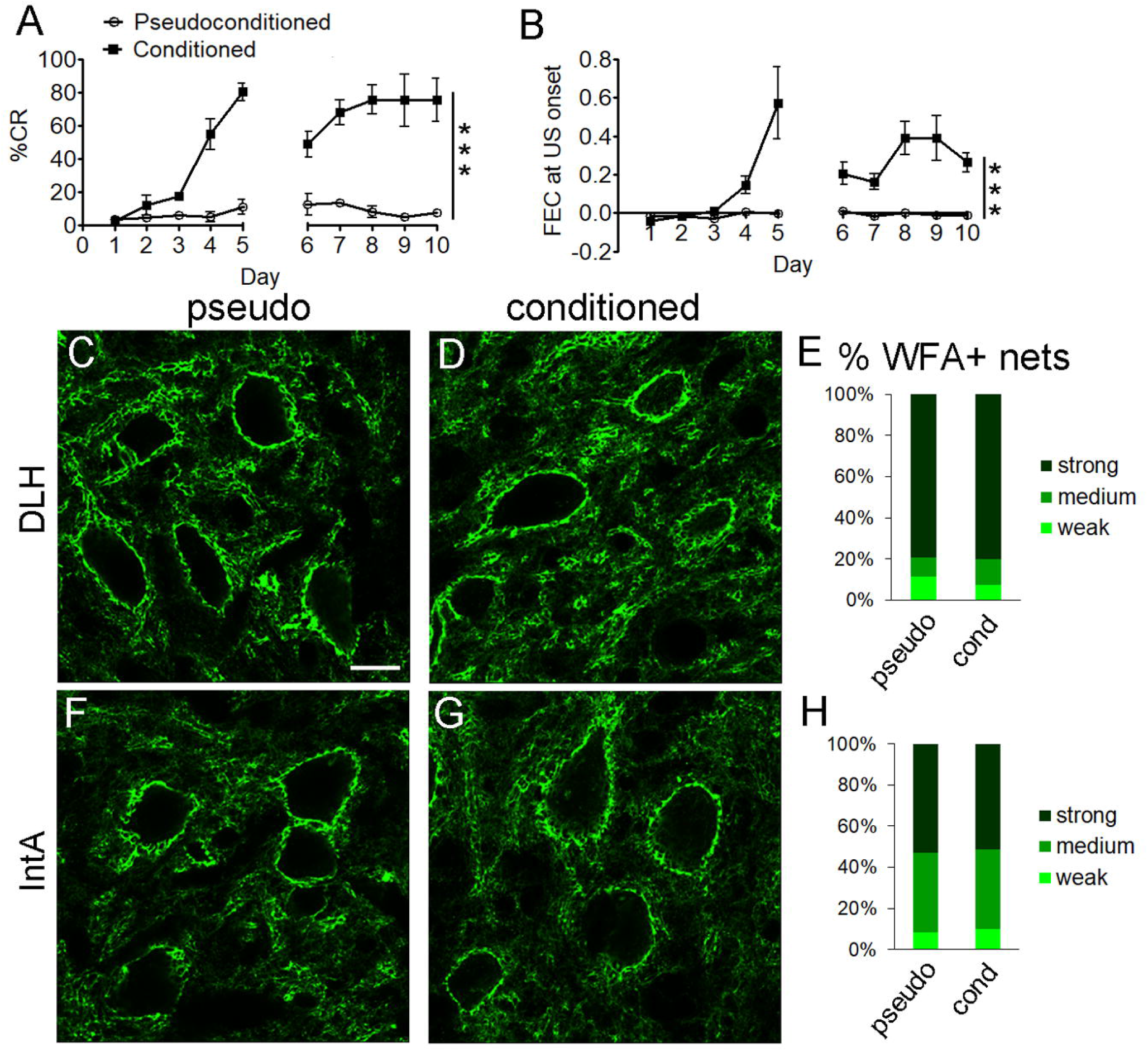
PNNs are restored during consolidation of EBC memory. (A, B) Mice were subjected to 10 sessions of EBC (conditioned) or pseudoconditioned stimulation. Conditioned mice learn during 10 days of training, as shown by their %CR (A) and FEC at the US onset (B). (C-H) WFA staining intensity of PNNs in the DLH (C-E) and the lateral IntA (F-H) is not different between conditioned mice and pseudoconditioned (pseudo) mice on day 10. The frequency distribution of weak, medium, strong WFA+ nets in the DLH and IntA is shown in (E) and (H), respectively. IntA = anterior interpositus, DLH = dorsolateral hump. Scale bar: 15 um in C (also applies to D, F, G). *** P < 0.001.

### PNN digestion enhances acquisition, but impairs retention of eyeblink conditioning

Based on the observed transient modulation of PNNs during EBC, we asked whether PNNs are necessary for acquisition and/or retention of this associative memory. To address this question, we digested PNN CSPGs in the IntA and DLH with ch’ase overexpressed by means of a lentiviral vector (LV) to ensure long-term, stable expression of the enzyme (Bosch et al., 2012, Burnside et al., 2018). Control mice received injections of LV overexpressing GFP. In LV-GFP mice, GFP expression was detected in the DCN (Fig. 3A) as early as one week post-injection and this was maintained at least up to 7 weeks (longest time point analyzed). CSPG amount was assessed by WFA labeling. In LV-ch’ase mice, WFA staining in the IntA and DLH was virtually completely abolished (~95% decrease; Fig. 3B-D). Qualitative observations revealed that the extent of PNN digestion was comparable in animals sacrificed at 2, 4 and 7 weeks after LV-ch’ase injection (not shown).

**Figure 3.**
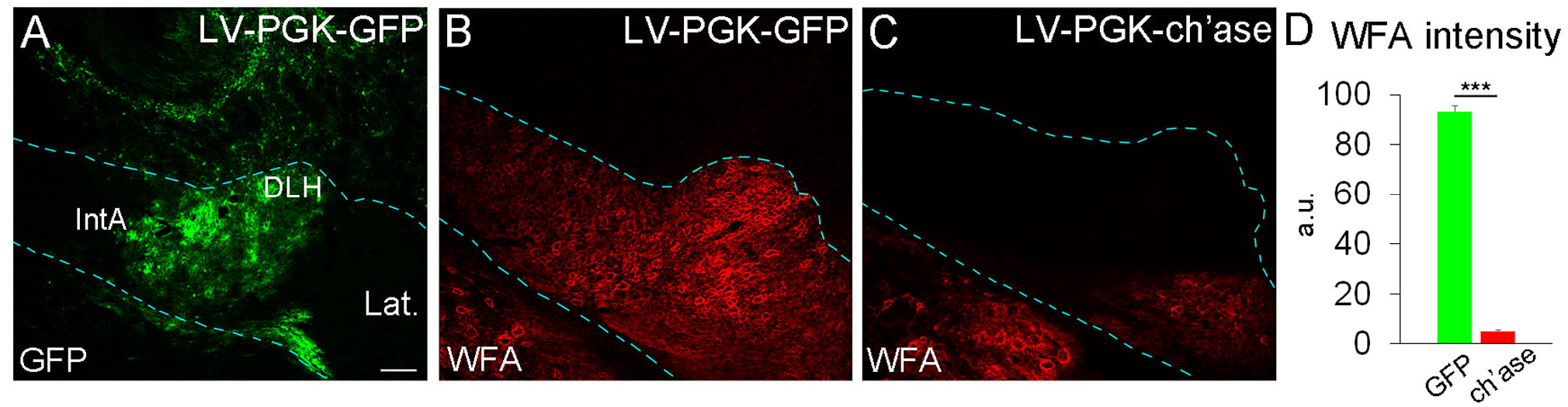
Transduction efficiency of LV-PGK-GFP and LV-PGK-ch’ase in the DCN. (A) Four weeks after LV-GFP injection, GFP is strongly overexpressed in the DCN and the cerebellar cortex (dorsal to the DCN). (B, C) Perineuronal nets, detected by WFA, are dramatically reduced following LV-ch’ase injection in the DCN (C), when compared to DCN injected with LV-GFP (B). D shows the quantification of WFA intensity in the IntA of GFP- and ch’ase-injected mice. In A-C, the DCN are outlined by dashed lines. IntA = anterior interpositus, DLH = dorsolateral hump, Lat. = lateral nucleus. Scale bar: 100 um in A (also applies to B, C). *** P < 0.001.

First, we examined whether removal of PNN-CSPGs by LV-ch’ase affected learning rates. Both LV-GFP (N=22) and LV-ch’ase (N=19) mice showed a significant increase in % of CRs over the course of EBC acquisition [LV-GFP: from 4% to 50%; LV-ch’ase: from 6% to 70%; F(4,39.447)=56.613, P<0.001; Fig. 4B]. However, LV-ch’ase mice learned significantly faster and better [group: F(1,41.530)=6.672, P=0.013; interaction: F(4,39.447)=3.403, P=0.018; Fig. 4B]. This effect was supported by an accelerated increase in FEC at US onset [day: F(4,39.457)=30.832, P<0.001; group: F(1,41.209)=4.691, P=0.036; interaction: F(4,39.457)=2.074, P=0.103, GLMM; Fig. 4C]. These results indicate that ch’ase overexpression in the IntA and DLH improves cerebellum-dependent learning.

**Figure 4.**
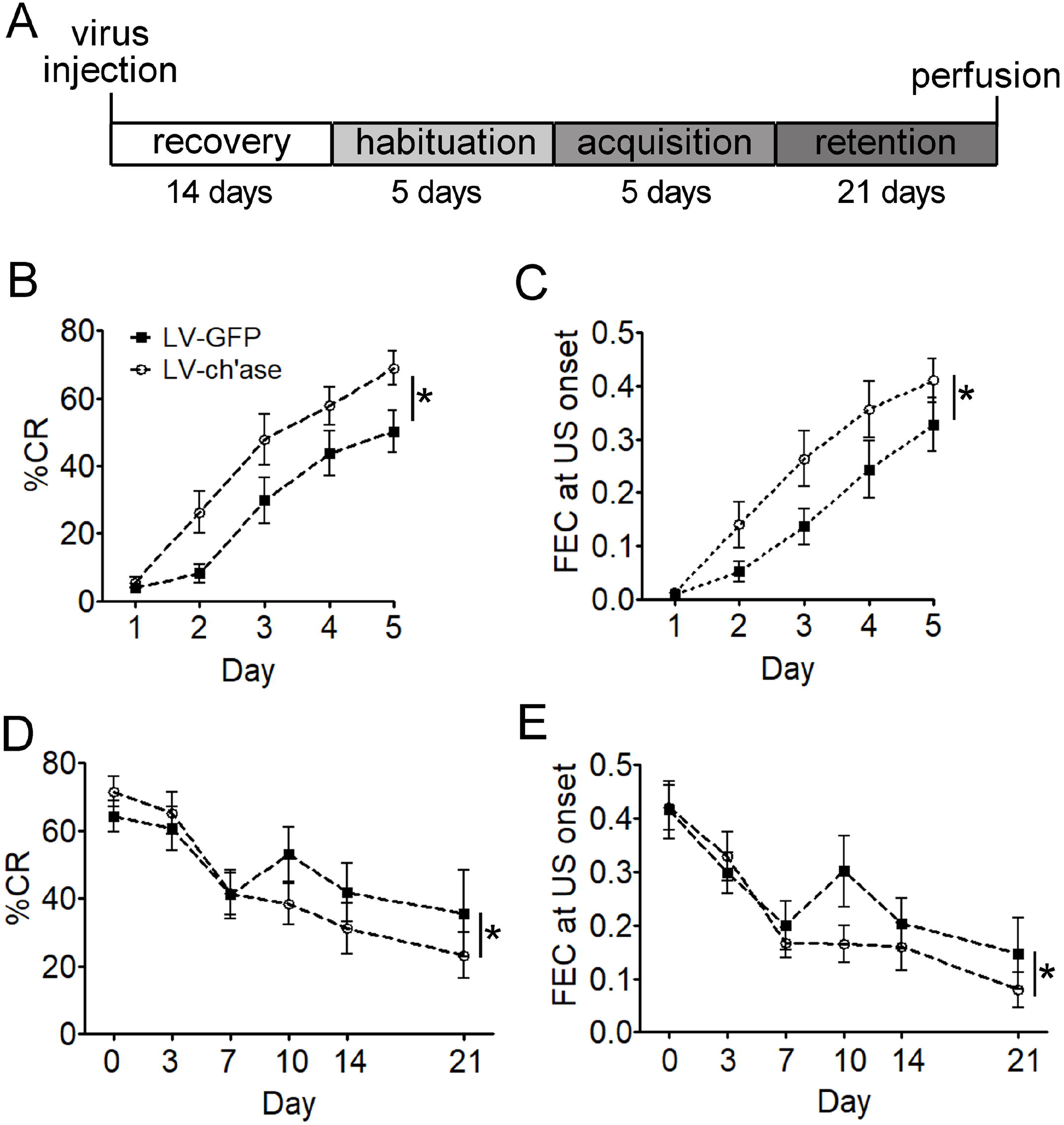
Effect of PNN digestion injections on EBC acquisition and retention. (A) Timeline showing the experimental design and duration of every phase of EBC. Days between phases are not shown. (B-E) EBC performance was calculated as %CR and FEC at US onset, and was compared between LV-GFP and LV-ch’ase mice during memory acquisition (B, C) and memory retention (D, E). During memory acquisition we found an increase in %CR (B) and FEC at US onset (C) over 5 days in both groups, as well as a significant difference between groups and an interaction effect, with ch’ase-mice showing better performance. During memory retention ch’ase-mice show worse performance, with %CR significantly different in the interaction days x groups (D) and FEC at US onset significantly different between groups from retention day 10 onwards (E). * P < 0.05.

Since we observed that WFA staining intensity was restored after 10 days of EBC (Fig. 2C-H), we hypothesized that long-term removal of PNNs would disrupt EBC retention. To test this hypothesis, we compared EBC performance over a period of 21 days, during which mice were intermittently retrained with short training sessions. Memory retention sessions consisted of 25% of a full acquisition session to reliably measure performance while attempting to retain a balance between memory reacquisition and extinction. For this test we included only mice that had shown evidence of learning during the acquisition phase, *i.e*. mice reaching >35% CR trials on day 5 of acquisition (LV-GFP: N=16; LV-ch’ase: N=18). Performance metrics of these mice on acquisition day 5 were comparable [LV-GFP versus LV-ch’ase: %CR: 64.2 ± 18.6% versus 71.4 ± 18.9%, t(32)=-1.124, P=0.27; FEC at US onset: 0.42 ± 0.22 versus 0.42 ± 0.18, t(32)=-0.075, P=0.941, two-tailed independent t-test]. We found that over the course of the memory retention phase, the performance of control mice decreased in the first week and then remained relatively stable, whereas LV-ch’ase mice exhibited a continuous decline with time, as shown by their % of CR trials [day: F(4,22.384)=11.533, P<0.001; group: F(1,30.703)=0.986, P=0.329; interaction: F(4,22.384)=3.185, P=0.033, GLMM; Fig. 4D] and FEC at US onset [D10-D21: day: F(2,23.956)=2.968; P=0.071; group: F(1,30.506)=4.339, P=0.046; interaction: F(2,23.956)=0.84, P=0.444, GLMM; Fig. 4E]. Together, these data support the hypothesis that retention of the EBC memory trace requires an intact PNN.

### Digestion of PNNs leads to an increased number of GABAergic terminals in the DCN

To pinpoint the neural substrate of the altered learning and memory capacities of mice after PNN digestion, we examined whether ch’ase alters the organization of IntA circuitry *in vivo*, focusing on morphological changes of GABAergic and glutamatergic axon terminals. The vast majority of synaptic terminals in the DCN are GABAergic and belong to Purkinje cells, as revealed by double staining for VGAT (which is contained in GABAergic terminals) and calbindin (which is expressed by Purkinje cells; Fig. S2A, B; De Zeeuw and Berrebi, 1995). GABAergic terminals are particularly abundant around the soma of DCN neurons (Fig. S2A). PNNs in the DCN surround excitatory neurons (Carulli et al., 2006; Hirono et al., 2018), which are characterized by a larger cell body size than that of inhibitory neurons (Uusisaari et al., 2007). Therefore, the effect of ch’ase on the morphology of GABAergic terminals was evaluated around the soma of large neurons (cell body size >250 um^2^, Uusisaari et al., 2007). We found that in ch’ase treated cerebella, GABAergic terminals appeared to be less distinct, forming a more continuous layer around the neuronal soma than in control cerebella (Fig. 5A-H). Indeed, the number of VGAT-negative spaces along the neuronal soma in ch’ase-injected IntA was strongly decreased when compared to the uninjected side or GFP-injected IntA (uninjected: 185.44 ± 7.20 troughs in VGAT intensity profile/mm neuronal membrane; n = 34 neurons ; GFP: 161.29 ± 6.24; n = 52 neurons; ch’ase: 86.24 ± 7.33, n = 33 neurons; one-way ANOVA F(2,116) = 42.24; P<0.001; Tukey’s post-hoc test: uninjected or GFP versus ch’ase, P<0.001; Fig. S3A-D). No difference between GFP-injected and uninjected IntA was detected (Tukey’s post-hoc test, P>0.05; Fig. S3A, B, D).

**Figure 5.**
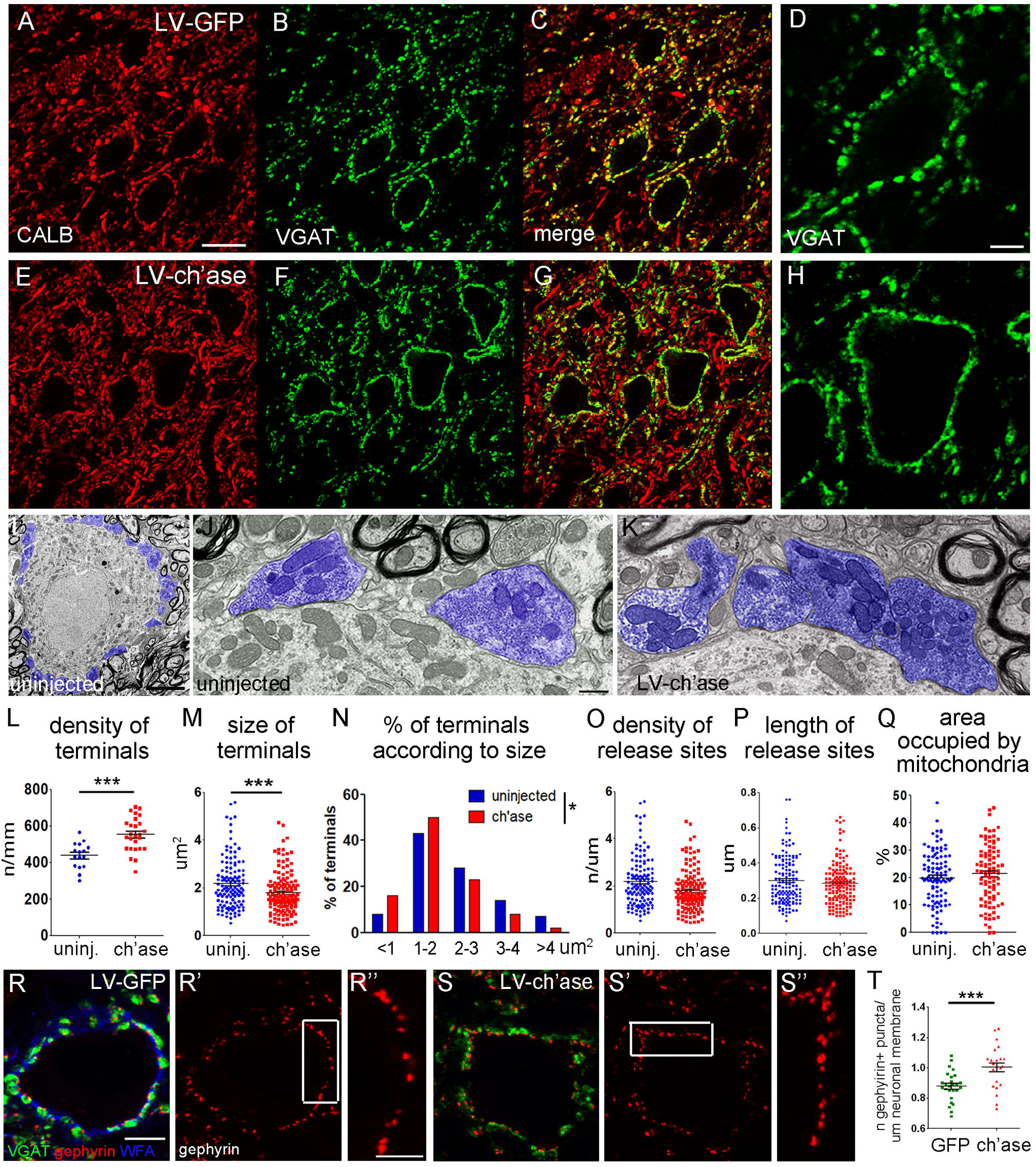
Effect of ch’ase on plasticity of GABAergic terminals in the DCN. (A-H) Immunolabeling for VGAT and calbindin (CALB) in the IntA of LV-PGK-GFP (A-D) and LV-PGK-ch’ase (E-H) injected mice at 4 weeks after virus injection. GABAergic terminals, including Purkinje cell boutons, appear as discrete puncta (abundant around DCN neuronal somata) in GFP-injected mice (A-D). See also Fig. S2. After ch’ase they show a more continuous distribution (E-H). See also Fig. S3. (I) Representative electronmicrograph of a control DCN neuron (no LV-ch’ase injection), surrounded by GABAergic terminals (digitally blue color coded). See also Fig. S4. (J) GABAergic terminals (blue) in uninjected DCN. (K) GABAergic terminals (blue) in ch’ase-injected DCN. (L-Q) Quantitations in EM pictures. (L) Density (number/mm neuronal membrane) of GABAergic terminals. Segments of neuronal membrane are shown as individual elements. (M) Size of GABAergic terminals (individual terminals are shown). (N) Frequency distribution of terminals according to size. (O) Number of release sites/um of membrane of synaptic terminal adjacent to the post-synaptic neuron (individual terminals are shown). (P) Length of individual release sites. (Q) Fraction of the area occupied by mitochondria in individual GABA+ synaptic terminals. (R-S’’) Immunolabeling with anti-gephyrin antibodies (red) shows gephyrin+ clusters juxtaposed to GABAergic terminals (which were stained by anti-VGAT antibodies, green) around DCN neurons in LV-GFP (R-R’’) and LV-chase-mice (S-S’’). WFA (blue) reveals the PNN around a control neuron (R) but is not visible around a ch’ase treated neuron (S). (T) Number of gephyrin+ clusters/um neuronal membrane (individual membrane segments are shown). Scale bar: 20 um in A (also applies to B, C, E-G); 5 um in D (also applies to H) and I; 0.5 um in J (also applies to K); 5 um in R (also applies to R’, S, S’); 2.5 um in R” (also applies to S’’). uninj. = uninjected. * P < 0.05, *** P < 0.001.

To study the ultrastructure of the GABAergic synapses following ch’ase-treatment, we performed electron microscopy. GABAergic boutons were identified by the presence of oval synaptic vesicles and symmetric release sites. Glutamatergic terminals were identified by the presence of rounded synaptic vesicles and thick post-synaptic densities (Uchizono, 1965; Kleim et al., 1998; Kleim et al., 2002; Fig. S4A-B’). In accordance with the immunohistochemical data, GABAergic boutons were abundant around the cell body of DCN neurons (Fig. 5I). After ch’ase treatment, GABAergic boutons did not display gross morphological abnormalities, showing clear synaptic vesicles, mitochondria and release sites (Fig. 5K). However, ch’ase induced a substantial increase in the number of GABA+ terminals (uninjected: 438.62 ± 17.75 terminals/mm; ch’ase: 553.47 ± 18.27 terminals/mm; Student’s t-test t_40_ = 4.23, P<0.001; Fig. 5J-L). In some cases, in ch’ase-treated cerebella, inhibitory terminals appeared “squeezed” along the neuronal membrane (Fig. 5K). Indeed, the average distance between GABAergic terminals was much lower than in control DCN (97.44 ± 14.66 nm versus 183.05 ± 27.64 nm, Student’s t-test t_60_ = 3.01, P<0.01). Moreover, the average size of GABAergic terminals was decreased after ch’ase (uninjected: 2.19 ± 0.096 um^2^; ch’ase: 1.79 ± 0.07 um^2^; Student’s t-test t_229_ = 3.34, P=0.001; Fig. 5M), due to an increased percentage of terminals with a small size (X^2^_4_ = 9.52, P< 0.05; Fig. 5N). The number of release sites/bouton was not different between the uninjected and ch’ase-injected DCN (uninjected: 1.05 ± 0.069 n/um; ch’ase: 1.17 ± 0.060 n/um; Student’s t-test t_147_ = 1.33, P=0.18; Fig. 5O), suggesting that ch’ase elicits the formation of new GABAergic terminals in the DCN, which are endowed with release sites and, thus, may be functional. No change in the length of release sites was found (uninjected: 300.00 ± 11.19 nm; ch’ase: 283.68 ± 9.61 nm; Student’s t-test t_296_ = 1.11, P=0.27; Fig. 5P). Synaptic mitochondria play a crucial role in the maintenance of homeostasis of presynaptic terminals and can divide, fuse and redistribute within the cell in response to various physiological cues (Vos et al., 2010). To assess whether ch’ase affects the amount of mitochondria in GABAergic terminals, we evaluated the percentage of area occupied by mitochondria in each terminal. No difference was detected between uninjected and ch’ase-side (uninjected: 19.84% ± 0.98%; ch’ase: 21.33% ± 1.14%; Student’s t-test t_191_ = 1.00, P=0.32; Fig. 5Q).

To further support the hypothesis that newly formed GABAergic pre-synaptic elements may be functional, we evaluated the number of gephyrin clusters using immunocytochemistry at the light microscopic level. Gephyrin is a scaffold protein that anchors GABA_A_ receptors at the post-synaptic membrane of inhibitory synapses, and is essential for the formation and stability of the GABAergic synapse (Tyagarajan and Fritschy, 2014). We found that the number of gephyrin clusters was higher in ch’ase mice than in control mice (LV-GFP: 0.88 ± 0.02 puncta/um neuronal membrane, n = 26 neurons; ch’ase: 1.00 ± 0.03, n = 23 neurons; Student’s t-test t_47_ = 3.69, P<0.001; Fig. 5R-T), confirming that following PNN digestion new, functional GABAergic synapses have been formed.

### Digestion of PNNs leads to a decreased number of glutamatergic terminals

We next evaluated whether ch’ase affects the number and size of glutamatergic terminals. Glutamatergic terminals in the DCN, including the IntA, belong to collaterals of mossy fibers and inferior olive axons, which mainly contact the dendritic compartment of DCN neurons (De Zeeuw and Berrebi, 1995). Since mossy fiber terminals contain VGLUT1 and/or VGLUT2, and olivary axon terminals contain VGLUT2 (Hioki et al., 2003; Mao et al., 2018), we visualized glutamatergic terminals by means of anti-VGLUT1 and anti-VGLUT2 antibodies. In control mice, VGLUT1+ and VGLUT2+ puncta were scattered throughout the IntA, with VGLUT1+ terminals being more numerous than VGLUT2+ terminals (Fig. 6A-C, G, I; Mao et al., 2018). Four weeks following LV-ch’ase injection, the number of VGLUT1+ terminals was significantly reduced (GFP: 35301.51 ± 961.91 terminals/mm^2^; ch’ase: 30342.28 ± 1011.63; Student’s t-test t_12_ = 3.55, P<0.01; Fig. 6A, C, D, F, G), whereas the number of VGLUT2+ terminals did not significantly change (GFP: 27969.37 ± 1471.89 terminals/mm^2^; ch’ase: 29772.64 ± 1591.36; Student’s t-test t_12_ = 0.83, P=0.42; Fig. 6B, C, E, F, I). The size of VGLUT1+ and VGLUT2+ terminals did not change either (VGLUT1: GFP, 1.18 ± 0.034 um^2^; ch’ase, 1.16 ± 0.042 um^2^; Student’s t-test t_12_ = 0.42, P=0.68; Fig. 6H; VGLUT2: GFP, 1.26 ± 0.019 um^2^; ch’ase, 1.30 ± 0.011 um^2^; Student’s t-test t_12_ = 1.36, P=0.20; Fig. 6J).

**Figure 6.**
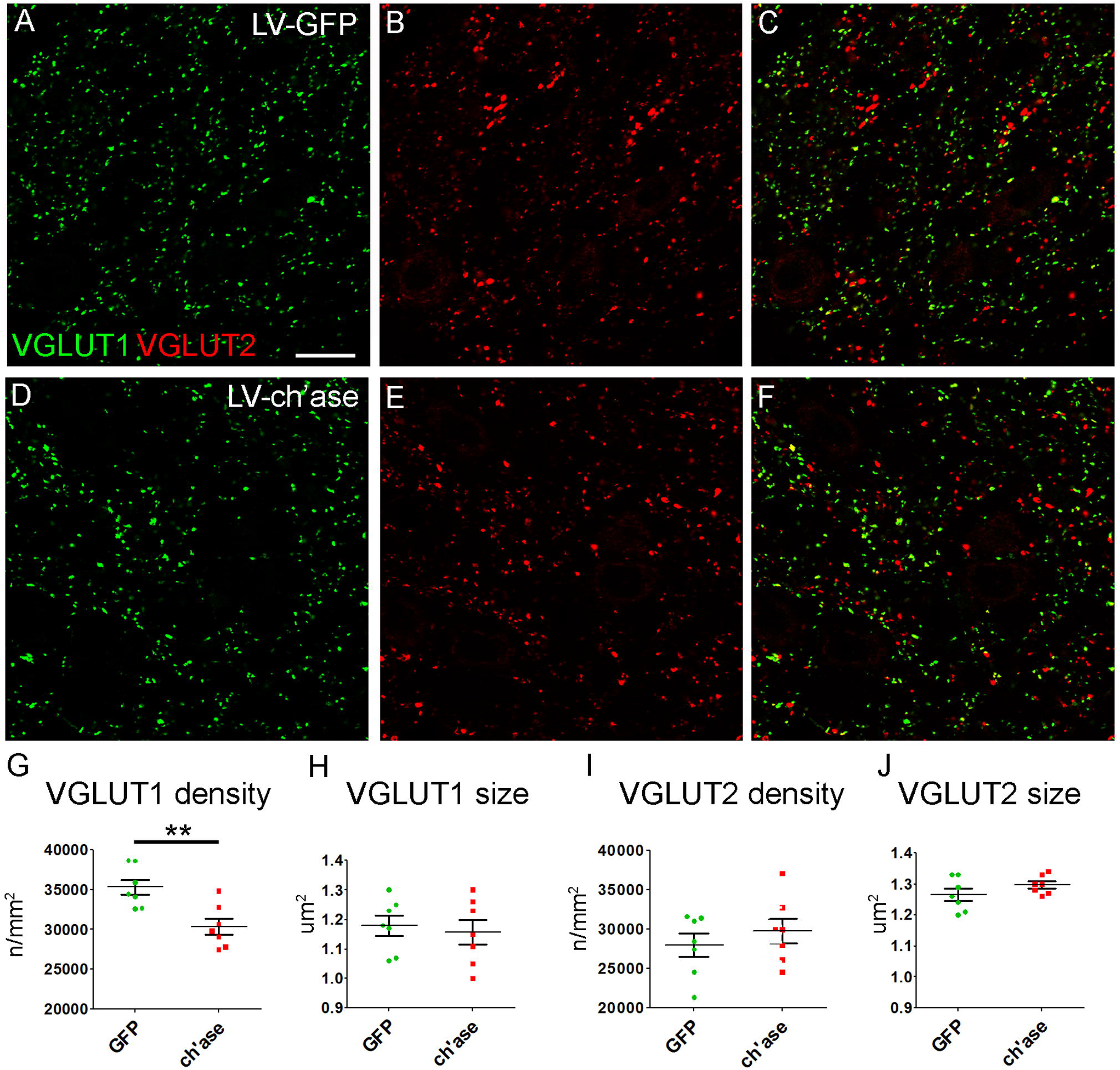
Effect of ch’ase on plasticity of glutamatergic terminals in the DCN. (A-F) Immunolabeling for VGLUT1 (green) and VGLUT2 (red) in the IntA of LV-PGK-GFP (A-C) and LV-PGK-ch’ase (D-F) injected mice at 4 weeks after virus injection. The number of VGLUT1+ terminals/mm^2^ (density) is significantly reduced after ch’ase (G), whereas VGLUT2 density remains unchanged (I). See also Fig. S5. The size of VGLUT1+ (H) and VGLUT2+ terminals (J) is not different between control and ch’ase-mice. Scale bar: 25 um in A (also applies to B-F). ** P < 0.01.

An increase in the number of mossy fiber varicosities in the DLH has been reported in association with learning during EBC, pointing to a role of synaptogenesis of mossy fibers in the formation of EBC memory (Boele et al., 2013). To examine whether the group difference in consolidation of the memory trace observed after ch’ase may be related to a loss of mossy fiber synapses induced by PNN digestion, we examined the number of VGLUT1+ terminals in the DLH of LV-GFP and LV-ch’ase mice at the end of EBC retention phase. We found a significant decrease in VGLUT1 density in conditioned ch’ase-treated mice when compared to conditioned GFP-mice (GFP, 46221.63 ± 625.74; ch’ase, 41387.06 ± 1516.97; Student’s t-test t_6_ = 2.95, P<0.05; Fig. S5A-C). Overall, PNN digestion *in vivo* triggers remarkable plasticity of synaptic connections in the DCN, namely an increase in the number of inhibitory synapses and a decrease in the number of excitatory terminals.

### PNN digestion alters electrophysiological properties of DCN neurons in awake mice

To evaluate the role of PNNs in the regulation of spike activity of DCN neurons *in vivo*, we made extracellular single unit recordings targeted at the lateral IntA and DLH in awake behaving mice. Recordings were obtained from LV-GFP (N=35 neurons in 6 mice) and LV-ch’ase mice (N=38 neurons in 5 mice). Recording location was estimated *post-hoc* after visualization of neurobiotin, which was added to the pipette solution and pressure-injected at the recording site. Neurobiotin staining appeared prominent in the IntA, while diffusing gradually into directly surrounding areas (Fig. S6A-C), confirming that our recordings were targeted at the correct region. Furthermore, we only included neurons that had been recorded in the IntA region with fully digested PNNs, as evaluated by *post-hoc* WFA staining. Spontaneous spike activity was recorded in awake mice during periods of quiet wakefulness, *i.e*. no locomotion or obvious movements occurred and no stimuli were presented. We found that the baseline firing frequency of recorded neurons in LV-ch’ase mice was significantly lower than that in LV-GFP mice (LV-GFP: 53.8 ± 31.4 Hz; LV-ch’ase: 35.6 ± 31.3 Hz; MWU-test U = 465, P=0.027; Fig. 7A, B). No difference in coefficient of variation (CV) of spiking (LV-GFP: 0.58 ± 0.25; LV-ch’ase: 0.80 ± 0.60; MWU-test U = 730, P=0.343; Fig. 7C) or average CV of two adjacent inter-spike intervals (CV2; LV-GFP: 0.50 ± 0.14; LV-ch’ase: 0.54 ± 0.21; Student’s t-test t_64.86_ = −0.821, P=0.415; Fig. 7D) was found between groups. These findings demonstrate that ch’ase-induced digestion of PNNs in the IntA leads to lower neuronal baseline spike activity *in vivo*.

**Figure 7.**
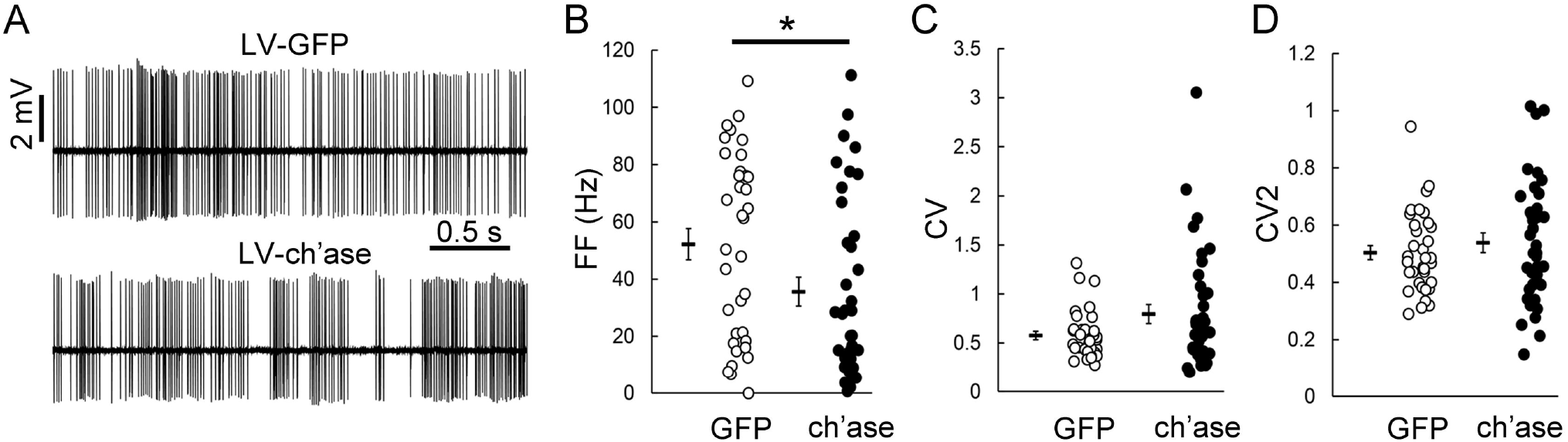
Spontaneous activity of IntA neurons after PNN digestion is reduced. (A) Representative traces of extracellular neuronal recordings in awake behaving LV-GFP (upper trace) and LV-ch’ase mice (lower trace). The firing frequency (FF) of IntA neurons is significantly reduced in LV-ch’ase mice when compared to LV-GFP mice (B), whereas CV (C) and CV2 values (D) are comparable. See Fig. S6 for recordings location. * P < 0.05.

## Discussion

In this study we have demonstrated that: i) PNNs in the DCN are reduced during EBC memory acquisition, but are restored when memories are consolidated; ii) digestion of PNNs in the DCN increases and accelerates EBC learning rate, but impairs memory retention; iii) PNN digestion causes substantial remodeling of DCN connections, with an increase in inhibitory synapses and a decrease in excitatory synapses; and iv) PNN digestion induces a reduction in the spontaneous firing activity of DCN neurons *in vivo*. Overall, we show that PNN modulation is a critical factor for dynamic control of DCN connectivity and, consequently, cerebellum-dependent learning (Fig. 8).

**Figure 8.**
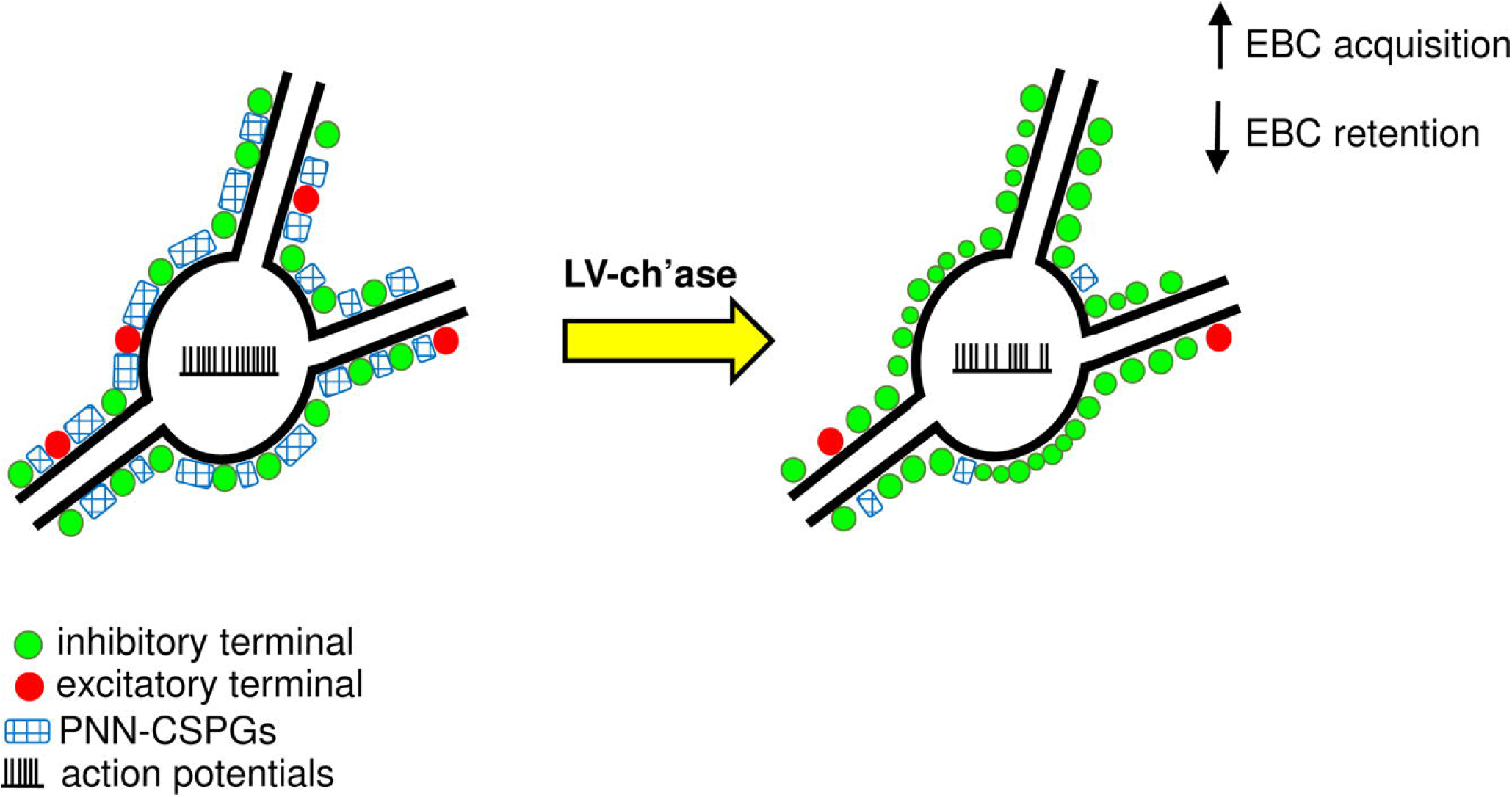
Schematic view of anatomical, electrophysiological and behavioral changes induced by PNN digestion in the DCN. The scheme shows that ch’ase overexpression in the DCN induces robust PNN-CSPG digestion around DCN neurons, which leads to structural synaptic remodeling (increased inhibitory terminals and decreased excitatory terminals) and decreased neuronal firing frequency, thereby impacting the formation and storage of associative memories.

### PNN dynamics and EBC

We found that PNNs exhibit very dynamic changes during acquisition and consolidation of cerebellum-dependent associative memories. Acquisition of EBC memories was accompanied by a decrease in the expression of PNN-CSPGs in DCN areas that are involved in the control of this type of learning, whereas, after consolidation of EBC memories, PNN-CSPG levels were not different from the control situation, suggestive of PNNs reinstatement. CSPGs are known to inhibit axonal growth (Kwok et al. 2008). Moreover, molecules interacting with CSPGs in the nets, such as Semaphorin3A (Sema3A) and Otx2, inhibit plasticity (Beurdeley et al., 201;, Spatazza et al., 2013; Boggio et al., 2019). Sema3A is also present in the DCN and Sema3A downregulation is associated with axonal remodeling therein (Carulli et al., 2013). Thus, we propose that reduced levels of PNN-CSPGs during EBC acquisition may allow synaptic remodelling, which, in turn, may drive the learning process. Indeed, an increased number of Purkinje cell synapses have been observed in the IntA of EBC trained mice (personal observations Broersen, Carulli, Canto, De Zeeuw). On the other hand, restored expression levels of CSPGs might promote synapse stabilization (Geissler et al., 2013), which is necessary for the subsequent maintenance of memory traces (Kandel et al., 2014).

Expression of PNNs in the brain, including the DCN, has been previously shown to be reduced following enriched environmental stimulation (Sale et al., 2007, Foscarin et al., 2011), locomotion training (Smith et al., 2015), injury (Carmichael et al., 2005; Carulli et al., 2013; Faralli et al., 2016) or drug exposure (Van den Oever et al., 2010; Vazquez-Sanroman et al., 2015; Slaker et al., 2018). Our study is the first to show a change in PNN expression during learning of new associations. In the context of learning and memory, dynamic regulation of PNNs around Golgi neurons in the cerebellar cortex has also been shown to be an important component of drug memories retention (Carbo-Gas et al., 2017). Moreover, PNNs in the auditory cortex are increased after fear conditioning (Banerjee et al., 2017). Thus, while elaborating on the work of others, our study is the first to indicate that the expression of PNNs can be up- and down-regulated during the acquisition and consolidation of newly formed associations, respectively.

Decreased expression of PNN components can occur by cleaving them with proteases, such as matrix metalloproteinases (MMPs) and ADAMTSs (A Disintegrin and Metalloprotease with thrombospondin motifs) (Tortorella et al., 1999; Nakamura et al., 2000; Levy et al., 2015; Senkov et al., 2014). In the cerebellum, MMP-9 mediates plasticity events induced by enriched environmental stimulation and contributes to a decrease in PNNs (Stamenkovic et al., 2017). Therefore, EBC may induce a finely-tuned, time-dependent modulation of such enzymes which, in turn, may control PNN stability. Moreover, because enriched environment as well as fear conditioning induces changes in the mRNA levels for PNN components (Foscarin et al., 2011; Banerjee et al., 2017), EBC may well exert a similar effect on the synthetic machinery of PNN molecules.

### PNN digestion and behavioural impact

Our observations of a dynamic regulation of PNN-CSPGs during EBC prompted us to examine whether PNN changes play a causative role in EBC learning and memory. Therefore, we enzymatically digested PNN-CSPGs in the DCN by overexpressing ch’ase in a lentiviral vector. The *ch’ase* gene, which is expressed in its original format in bacteria, was modified to allow efficient secretion of active chondroitinase from mammalian cells (Muir et al., 2010). The viral approach ensures a long-term expression of the enzyme in the cortex and spinal cord (Zhao et al., 2011; Bartus et al., 2014). In the cerebellum, we observed strong PNN-CSPG digestion in the DCN, which resulted in substantial acceleration and amelioration of learning. The direct injection of the enzyme into the DCN has been shown to have a smaller effect, inducing a partial digestion of CSPGs and an increase in the learning rate only at late stages of the acquisition phase (Hirono et al., 2018). Thus, removal of PNN-CSPGs with our lentiviral approach induced faster and better associative motor learning, allowing us to establish a causal role for a decrease in PNN during acquisition. The impact of PNN digestion with ch’ase on learning has also been studied in the context of cognitive or emotional processes. For example, digestion of PNNs in the auditory cortex results in enhanced performance in animals trained in a cue reversal learning task (Happel et al., 2014). Likewise, degradation of PNNs in the amygdala can induce erasure of fear memories or drug memories via extinction, which is a form of learning (Gogolla et al., 2009; Xue et al., 2014). Importantly, our study shows for the first time that digestion of PNN-CSPGs causes a decrease in EBC retention over time, suggesting that PNNs are implicated in the consolidation of previously acquired associative motor memories. Although PNN digestion in cortical areas negatively affects the storage of drug-associated memories (Slaker et al., 2015) and fear memories (Banerjee et al. 2017; Thompson et al. 2018), it has opposite effects on object recognition memory. This suggests that PNNs may play different roles in distinct types of memory (e.g. working memories versus associative memories), depending on the organisation and timing-dependent properties of the underlying neuronal network involved.

It is interesting to speculate on a role of DCN PNNs in the control of a critical period for retention of associative motor memories regulated by the cerebellum. Retention of EBC memories in rats is labile when memories are acquired at young age (postnatal day 17), but not when they are acquired in adulthood (Nicholson et al., 2003; Brown and Freeman, 2014, 2016). Notably, PNNs in the DCN are still very immature between P14 and P21 (Carulli et al., 2007), raising the possibility that memory consolidation in the DCN is subject to a critical period during early postnatal development, which depends on PNN maturation.

### PNNs and DCN synaptic connectivity

Our study is the first to show that synaptic connectivity in the DCN is profoundly altered when PNN-CSPGs are digested, as we observed a shift of the excitatory/inhibitory balance towards an increased inhibition. Ch’ase induced a strong increase in the number of Purkinje cell synapses on DCN neurons. Purkinje cells are essential for the control of EBC acquisition rate and timing (McCormick and Thompson, 1984; Perrett et al., 1993; Garcia et al., 1999; Christian and Thompson, 2003). Therefore, fine-regulating Purkinje cell input may facilitate learning. In contrast to our results, the study by Hirono et al. (2018) shows no structural modifications of Purkinje cell terminals in acute cerebellar slices treated with ch’ase. This discrepancy could be explained by a different efficacy of ch’ase [25% PNN reduction in Hirono et al. (2018) versus 90% PNN reduction in our study], different age of the animals (adolescent versus adult), and/or different model employed (*in vitro* versus *in vivo*). Overall, our data strongly support a role for PNN-CSPGs in restricting Purkinje axon growth and synapse formation. Likewise, the growth potential of Purkinje axon collaterals in the cerebellar cortex is under control of CSPGs that are diffusely distributed in the neuropil (Corvetti and Rossi, 2005).

We also found that ch’ase induced a decrease in the density of VGLUT1+ terminals in the dorsolateral hump (DHL) and adjacent areas of the anterior interposed nucleus. The observed differential effects of ch’ase on distinct types of cerebellar synapses might be due to a specific distribution of receptors for CSPGs (Duncan et al., 2019) and/or for Sema3A (Vo et al., 2013).

Pontine mossy fibers, which relay the CS to the cerebellum (Brodal et al., 1986; Knowlton and Thompson, 1988), express VGLUT1+ (Allen Brain Atlas). The pontine mossy fibers in the DHL sprout during paired eyeblink conditioning, but not during pseudo-conditioning with the same set of sensory inputs, and the number of sprouting fibers can be correlated to the amplitude of the conditioned response (Boele et al., 2013). We hypothesize that in ch’ase-treated mice mossy fiber synaptic contacts formed during the acquisition phase may, due to the absence of PNNs, not be stabilized, and, as a consequence, the memory trace cannot be consolidated. Thus, a net increase in the inhibitory tone onto DCN neurons may promote initially an enhanced learning rate, whereas a higher level of baseline excitation may be important for memory retention. How can this be reconciled with our observation that PNNs are also important for the baseline electrophysiological properties of DCN neurons? After ch’ase treatment, DCN neurons of awake behaving mice showed lower spontaneous firing frequencies. This change in the baseline firing rate of DCN neurons may be partly explained by the increased Purkinje cell innervation and/or the decreased density of excitatory terminals. Nonetheless, due to a presumed role of PNNs in preventing the free diffusion of potassium or sodium ions in the extracellular space (Hartig et al., 1999), we cannot exclude that a decreased reservoir of available cations following ch’ase administration also provides a more direct impact onto the firing frequency of DCN neurons. Importantly, reduced baseline activity of DCN neurons may affect the level of rebound firing (Aizenman and Linden, 1999; Zheng and Raman, 2009), which in turn may play a role in the induction of DCN plasticity and the strength of the CR (Kistler et al., 2000; Wetmore et al., 2008; De Zeeuw et al., 2011). Moreover, since the simple spike activity of Purkinje cells is temporarily suppressed during the actual expression of a CR (Rasmussen et al., 2008; Halverson et al., 2015; ten Brinke et al., 2015), we expect the level of rebound firing in the DCN during CR expression to be reduced when their baseline firing is lower (Ten Brinke et al., 2017).

### Conclusions

We provide compelling evidence that PNNs are indispensable for the control of specific connection patterns in the DCN and for regulating functional properties of DCN neurons. Patterns of not only modulatory, but also baseline activity of DCN neurons are essential for the formation and expression of associative memories. Since the cerebellum controls not only motor functions, but also social, cognitive and emotional functions (Sacchetti et al. 2002; Timmann et al. 2010; Wang et al. 2014; Gao et al., 2018), PNNs may be implicated in various cerebellum-dependent processes in both physiological and pathological conditions.

## Acknowledgements

The work was supported by University of Turin, La Maratò de TV3, International Foundation for Research in Paraplegia, the Netherlands Institute for Neuroscience, the Netherlands Organization for Scientific Research (NWO-ALW), the Dutch Organization for Medical Sciences (ZonMW), ERC-adv and ERC-POC. We are grateful to Bas Koekkoek and Ilja IJpelaar (Erasmus MC, Rotterdam, Netherlands) for their help with the eyeblink conditioning paradigm, Willemijn Ranzijn (Netherlands Institute for Neuroscience, Amsterdam, Netherlands) for her help with behavioral experiments and immunohistochemistry, Barbara Hobo, Cynthia Geelen and Anna Court (Netherlands Institute for Neuroscience, Amsterdam, Netherlands) for their excellent technical support.

## Author contributions

DC, JV, CIDZ, RB and CBC conceived and designed the study. DC performed surgery, immunohistochemistry, confocal acquisition, data analysis. RB performed behavioural experiments, electrophysiology and data analysis. FdW and EMM generated the viral vectors. HJB set up the eyeblink conditioning paradigm and equipments. MM performed behavioural experiments. MdW performed immunohistochemistry. SdV performed electron microscopy. DC, RB, CIDZ and JV wrote the paper with input from the other authors.

## Materials and methods

### Ethical statement

Adult male C57BL/6J mice (6-8 weeks old; Janvier labs) were socially housed with food and water *ad libitum*, in 12 h light and dark cycles. All procedures were approved by the animal committee of the Royal Dutch Academy of Arts & Sciences and adhered to the European guideline for the care and use of laboratory animals (Council Directive 86/6009/EEC).

### Eyeblink conditioning

For pedestal placement, mice were anesthetized with isoflurane (5% induction, 1.5-2% maintenance) and placed in a stereotaxic frame where their body temperature was kept at 37°C using a feedback-controlled heating pad. Eyes were covered with antimicrobial ointment to prevent drying (Terra-Cortril, Pfizer). A mid-sagittal incision of approximately 1 cm was made to expose the skull bone, on which primer (Optibond All-In-One, Kerr) was applied and treated with UV light. A 6 x 3 x 5.7 mm aluminum block (‘pedestal’) was attached to the skull bone with dental acrylic (Flowline, Hereaus Kulzer). Skin edges were attached to the dental acrylic using tissue glue (Histoacryl, Aesculap AG). Post-operative analgesia was given subcutaneously (Meloxicam, 0.4 mg/kg, Metacam, AUV). Mice were allowed to recover for at least two days following surgery prior to being subjected to behavioral analysis. Mice were subjected to eyeblink conditioning (EBC) using a camera setup or a setup based on a magnetic distance measurement technique (MDMT setup) (Koekkoek et al., 2002). In the MDMT setup mice were placed in light and sound-proof boxes, whereas in the camera setup mice were placed in a faraday cage with curtains to create darkness. In both cases mice were first habituated for 5 days with increasing duration (from 15 minutes on day 1 to 60 minutes on day 5). During habituations mice were head-fixed to a horizontal bar and could freely walk on a cylindrical treadmill in the dark to get accustomed to the environment (Chettih et al., 2011). For mice trained in the MDMT setup, under isoflurane anesthesia hairs under the left eye were removed on day 3. From day 4 onward a small neodymium magnet (1.5 × 0.7 × 0.5 mm) was placed under the left eyelid using superglue (Bison) to allow detection of eyelid movements, thanks to a magnet sensor placed above the magnet, which detects changes in magnetic field. Ten minutes after closing the box, trainings were initiated. In the camera setup mice were placed on the treadmill and before fixation a thin layer of waterproof black mascara was applied on the left whiskers to prevent infrared light (IR) reflection. For all mice, from day 4 onward a sequence of 10 CS only - 2 US only - 10 CS only trials was presented to make mice acquainted with the stimuli. US was a 10ms corneal air puff (35-40 psi) delivered through a p20 pipette tip positioned ~5 mm from the left eye. CS was a 260ms green (camera setup) or green/blue (MDMT setup) LED light positioned ~7 cm in front of both eyes (Fig. 1C). US onset was 250ms after CS onset, both stimuli co-terminated and stimuli were presented with an inter-trial-interval (ITI) time of 12 ± 2sec (camera setup) or 10 ± 2sec (MDMT setup). Trials were initiated only if the eye was more than ~75% opened. Eyelid movements in the camera set up were recorded using a 250fps camera (scA640-120 gc, Basler) and the mouse eye was illuminated by an IR-light. Data was sampled at 2441Hz and stimuli were triggered using TDT System 3 (Tucker Davis Technologies) and NI-PXI (National Instruments) processors. Eyelid movements in the MDMT set up were captured at 1000Hz using custom-written LabVIEW (National Instruments) software (Koekkoek et al., 2002).

To study the effect of 5 days of EBC on PNNs in the DCN, we randomly assigned mice to one of three groups: conditioned (N = 5), pseudoconditioned (N = 5) or control (N = 6). Control mice received a pedestal but remained in the cage for the duration of the experiment. Trainings for these experiments were done with the camera setup. Mice in the conditioned group received a sequence of 3 US only and 100 CS-US paired trials presented with an ITI of 12 ± 2sec, 3 times per day for 5 consecutive days, totaling 15 training sessions. Mice in the pseudo-random group received a sequence of 3 US only trials followed by a random sequence of 100 CS only and 100 US only trials 3 times a day for 5 days. Pseudo-random stimuli were presented with an ITI of 6 ± 2sec to match the duration of the training of the conditioned group. To study the effect of 10 days of EBC on PNNs in the DCN, mice were conditioned (N = 3) or pseudoconditioned (N = 3) for 10 days (5 days/week, for 2 weeks). To randomize trials we used the ‘randi’ function of MATLAB (MathWorks) and saved the order in one of three templates which we used in an alternating manner each session. Performance of conditioned and pseudoconditioned mice was measured as the percentage of trials showing conditioned responses (%CR) and the fraction eyelid closure (FEC; 1 is fully closed, 0 is fully opened) at the onset of the US.

### Virus production

The GFP or chondroitinase (ch’ase) genes were inserted into lentiviral vectors for long-term expression, containing the mouse phosphoglycerate kinase (PGK) promoter. The vector has been produced according to Zhao et al. (2011). Briefly, the GFP or chondroitinase transgene was subcloned into a transfer plasmid such that the transgene could be transcribed into a packageable RNA upon transfection into HEK293T cells along with non-recombining plasmids that express lentiviral and VSV-G genes (Naldini et al. 1996; Dull et al. 1998). Each transfer plasmid specifies a vector RNA which contains, from 5’ to 3’, the RU5 fragment of the long terminal repeat (LTR), HIV-1 packaging signal in a Gag gene fragment, Rev response element, central polypurine tract/central termination sequence, PGK promoter with the transgene, woodchuck hepatitis virus post-transcriptional regulatory element, and self-inactivating (SIN) 3’ LTR. The vector RNAs are essentially identical except for the polylinker and transgene. The lentiviral vector was created with a transfer plasmid derived from pRRL via the SIN-W-PGK vector, with different polylinkers which includes an additional polypurine tract, and vector particles were generated by cotransfection with this plus two other plasmids (Déglon et al. 2000). Viral particles were concentrated by ultracentrifugation and the viral particle-containing pellet was resuspended in 0.1 M phosphate-buffered saline pH 7.4 (PBS) or in Dulbecco’s modified Eagle’s medium (DMEM), and stored at −80°C until further use. Viral titer was obtained by a p24 antigen ELISA assay (Perkin Elmer). The titer of LV-PGK-GFP was 3 x 10^10^ transducing units, the titer of LV-PGK-ch’ase was 1 x 10^10^ transducing units.

### Virus injections

Mice were deeply anesthetized using isoflurane and eyes were protected from drying using antibacterial eye cream (Terra-Cortril, Pfizer). Body temperature was maintained at 37°C using a temperature-controlled heating pad. A 1 cm longitudinal skin and muscle incision was made at the level of the occipital bone. For behavioral and electrophysiological evaluations, bilateral injections were performed. Two small craniotomies were done to expose the IV-V cerebellar lobule of the right and the left hemispheres. One μl of virus (LV-PGK-GFP or LV-PGK-ch’ase; titer matched at a final titer of 1 x 10^10^ transducing units) was injected into each interpositus nucleus (−6.0 mm from Bregma, +-2.0 mm lateral from midline, 2.0 mm depth) by a quartz capillary pipette (30-40 μm tip diameter) connected to a Harvard injection pump (speed: 0.15 μl/min). The pipette was left in place for 2 minutes after the injection and was then slowly retracted. For morphometric assessments of synapses, unilateral injections (in the right DCN) were performed and animals were sacrificed 4 weeks later. For EBC experiments, after virus injection a continuation of the first incision was made in the skin covering the top of the skull and a pedestal was stereotaxically placed (see above). The head skin was sutured and animals were given post-operative analgesia (0.4 mg/kg Meloxicam, Metacam).

### Eyeblink conditioning on virus-injected mice

To study the effects of PNN digestion on EBC acquisition and retention, N = 22 mice were injected with LV-PGK-GFP and N = 19 mice were injected with LV-PGK-ch’ase. After surgery, mice recovered for 14 days before initiation of habituation sessions (described above). Behavioral tests were performed blind to eliminate experimenter bias. Starting on day 21 after injections, mice were trained once a day for 5 consecutive days using the MDMT setup (*acquisition phase*). Each session consisted of 20 blocks of 1 US only - 10 CS-US paired - 1 CS only trials presented with an ITI of 10 ± 2sec, totaling 20 US only, 20 CS only and 200 paired trials per session. Following acquisition training, memory retention (*retention phase*) was measured in mice that had showed robust learning during the acquisition phase (%CR >35% on the last acquisition day; LV-GFP: N = 16; LV-ch’ase: N = 18). All these mice were subjected to retention sessions on day 3, 7, 10 after the last acquisition day, and a number of them was also subjected to retention sessions on day 14 (LV-GFP: N = 10; LV-ch’ase: N = 11) and 21 (LV-GFP: N = 6; LV-ch’ase: N = 7). Retention sessions consisted of 5 blocks of 1 US only - 10 CS-US paired - 1 CS only trials presented, totaling 5 US only, 5 CS only and 50 paired trials. Mice were transcardially perfused the day after the last retention day. The timeline of the experiment is shown in Figure 4A.

### Eyeblink data analysis

Data acquired on the camera setup was analyzed using custom-written MATLAB (MathWorks) scripts. Data were filtered using a Gaussian 50 Hz lowpass filter and event-related data were gathered based on TTL timestamps indicating trial start and CS and US stimuli. Trials were marked as unstable and were excluded if values during baseline (first 500 ms of trial) exceeded 5x SD of signal of baseline. To correct for the measurement bias during squinting following a US, all amplitudes were calculated relative to 66.7% of the average UR amplitude. UR amplitude was calculated as average of maxima during the first 450ms following a US in US only trials (or paired trials if not available). CRs were defined as eyelid closures exceeding 10% of UR amplitude during 50-250 ms after CS onset. Data from the MDMT setup was similarly analyzed, although sessions were excluded if more than 75% of trials were invalid. The following parameters were calculated: i) CR percentage, calculated by dividing the number of CR trials with the total amount of CR-eligible trials, multiplied by 100; ii) fraction eyelid closure (FEC) at US onset, calculated by dividing the value of the baseline-normalized signal at US onset (750 ms after trial start) with the average UR amplitude.

### Histological procedures

Mice were anaesthetized with an overdose of pentobarbital (Nembutal) and transcardially perfused with 100 ml of 4% paraformaldehyde in 0.12 M phosphate buffer. Brains were postfixed overnight at 4°C, then cryoprotected in 0.12 M phosphate buffer containing 30% sucrose at 4°C, until they sank. Cerebella were cut on a cryostat into 25 μm-thick coronal or sagittal sections and collected in phosphate-buffered saline (PBS). Primary antibodies were incubated overnight at 4°C in PBS containing 0.25% Triton X-100, and 5% fetal calf serum. Primary antibodies were: mouse anti-NeuN (Millipore, 1:500), chicken anti-GFP (Aves Labs, Tigard, Oregon, USA, 1:700), mouse anti-calbindin (Swant, 1:1500), rabbit anti-VGAT (Synaptic Systems, 1:1000), rabbit anti-VGLUT1 (Synaptic Systems, 1:1000), guinea pig anti-VGLUT2 (Synaptic Systems, 1:1000), mouse anti-gephyrin (Synaptic Systems, 1:500), mouse anti-MAP2 (Chemicon, 1:250). To visualize PNNs, we used *Wisteria floribunda* agglutinin (WFA), which binds CSPG sugar chains and is therefore a general marker of PNNs (Hartig et al. 1992). Sections were incubated in biotinylated WFA (Vector, 20 μg/ml) for 2 h at room temperature. After incubation in primary antibody or WFA, sections were incubated for 1 h at room temperature with one of the following fluorophore-conjugated secondary antibodies or streptavidin: donkey anti-mouse Cy3 (ThermoFisher Scientific), donkey anti-mouse Alexa Fluor 647 (ThermoFisher Scientific goat anti-chicken Alexa Fluor 488 (ThermoFisher Scientific), donkey anti-guinea pig Cy3 (ThermoFisher Scientific), donkey anti-rabbit Alexa Fluor 647 (ThermoFisher Scientific), streptavidin-488 (ThermoFisher Scientific), streptavidin-Cy3 (Jackson Immunoresearch). For each immunohistochemical reaction, slices from all experimental conditions were processed together and incubation times kept constant. After processing, sections were mounted on microscope slides with Tris-glycerol supplemented with 10% Mowiol (Calbiochem).

Histological preparations were examined under a Leica SP5 confocal microscope. Confocal images were taken at a resolution of 1024×1024 dpi and 50 Hz speed. Lasers intensity, gain and offset were maintained constant in each analysis. Quantitative and morphometric evaluations were performed by using Image J software (see below) and were conducted by a blinded experimenter. Adobe Photoshop 6.0 (Adobe Systems, San Jose, CA) was used to adjust image contrast and assemble the final plates.

### Quantification of WFA intensity

Quantification of the staining intensity of PNNs in 5 days-conditioned (N = 5), 5 days-pseudoconditioned (N = 5) or control (N = 6) mice was performed in the DLH, in the most lateral part of the anterior interpositus (IntA) and in the lateral nucleus in coronal sections on both sides. Sections were double labelled with NeuN antibodies and WFA. Confocal images (1 um thick) of sections of the anterior DCN (6.00 to 6.24 mm caudal to Bregma) were acquired under a 63× objective. Two-three sections/mouse were analysed. Analysis of WFA staining intensity around neurons with a visible NeuN+ nucleus was performed by ImageJ, as described in Foscarin et al. (2011). Briefly, for each analysed neuron we measured the brightness intensity of WFA (range 0-255), by randomly selecting 15 pixels (approximately equidistant from each other) surrounding the neuronal soma and calculating their average intensity [DLH: CTR right side, number of neurons (n) = 106, CTR left side n = 96; pseudo right side n = 81, pseudo left side n = 65; conditioned right side n = 66, conditioned left side n = 87; IntA: CTR right side n = 99, CTR left side n = 95; pseudo right side n = 63, pseudo left side n = 75; conditioned right side n = 72, conditioned left side n = 88; lateral nucleus: CTR right side n = 110, CTR left side n = 104; pseudo right side n = 82, pseudo left side n = 86; conditioned right side n = 75, conditioned left side n = 94]. The background brightness, taken from a non-stained region of the cerebellar molecular layer, was subtracted from the brightness measurements. Each net was assigned to one of three categories of staining intensity, ranging from the lowest to the highest value of WFA intensity: weak = 0-33%, medium = 34-66%, strong = 67-100% of maximum staining intensity. The same analysis has been performed in the DLH and the IntA of 10 days-conditioned (N = 3) and 10 days-pseudoconditioned (N = 3) mice (DLH: pseudo n = 53; conditioned right side n = 28, conditioned left side n = 27; IntA: pseudo n = 36; conditioned right side n = 19, conditioned left side n = 22).

Quantification of PNN digestion after injection of LV-ch’ase (N = 3) or LV-GFP (N = 3) was performed by evaluating WFA intensity in a selected area of the IntA (80,000 um^2^) in 3 sections/mouse. The background brightness, taken from the molecular layer of a not-digested lobule, was subtracted from the measurements.

### Quantification of gaps between VGAT-positive boutons

Single 0.5 um thick-confocal images of the IntA in sagittal sections have been collected under a 63× objective with 2x zoom. On such images, the intensity of VGAT staining around large size, excitatory neurons (cell body size > 250 um^2^, Uusisaari et al., 2007), which are the neurons bearing a PNN (Carulli et al. 2006; Hirono et al. 2018), has been plotted by using the ImageJ function “surface plot”. In the plot, the density (number/mm neuronal membrane) of troughs with value < 40 arbitrary unit (a.u.), followed by peaks with value > 100 a.u. has been quantified per each analysed neuron (LV-GFP, n = 52 neurons from 5 mice; LV-ch’ase, n = 33 neurons from 5 mice).

### Quantification of gephyrin-positive clusters

Single 0.5 um thick-confocal images of sagittal sections containing the IntA have been collected under a 63× objective with 3x zoom. On such images the density of gephyrin-positive clusters (number/um neuronal membrane) has been evaluated around large size neurons by means of ImageJ (LV-PGK-GFP, n = 26 segments of neuronal membrane from 3 mice; LV-PGK-ch’ase, n = 23 segments of neuronal membrane from 4 mice).

### Quantification of glutamatergic terminals

We estimated the density and size of VGLUT1 and VGLUT2-positive glutamatergic axon terminals in LV-GFP (N = 7) and LV-ch’ase (N = 7) injected mice. At least three sagittal sections containing the IntA were selected for each animal. We estimated the density of VGLUT1-positive terminals in in LV-GFP (N = 4) and LV-ch’ase (N = 4) injected mice subjected to EBC and perfused on retention day 21. At least three coronal sections containing the DLH were selected for each animal. A 0.5 um-thick confocal image/section was captured under a 63x objective. The ‘‘analyze particle’’ function of ImageJ was used to estimate the density (number of terminals/mm^2^) and size of boutons in each image (area of the image = 15,117 um^2^), after selecting the automatic threshold and watershed segmentation. Only elements with a size > 0.4 um^2^ were included in the analysis, to discard staining background artifacts. DCN neuronal somata were weakly VGLUT2-simmunopositive and were excluded from the analysis.

### Electron microscopy

Mice that received unilateral injection of LV-PGK-ch’ase (N = 3) were perfused with 4% paraformaldehyde in 0.12 M phosphate buffer. Brains were postfixed overnight at 4°C, then cryoprotected in 0.12 M phosphate buffer containing 30% sucrose at 4°C. Cerebella were cut on a cryostat into 25 μm-thick sagittal sections and collected in phosphate-buffered saline (PBS). Slices were washed with sodium cacodylate buffer and postfixed for 20 minutes in 1% osmium tetroxide and 1.5% potassium ferricyanide in milliQ, dehydrated and embedded in epoxy resin. Sagitally cut ultrathin sections (70 nm) were contrasted with uranyl acetate and lead citrate and analyzed using a FEI Technai 12 electron microscope.

Analysis of GABAergic terminals in electron micicrographs has been performed on large size neurons (cell body size > 250 um^2^) of the IntA in 3 mice, which received LV-cha’se injection unilaterally. Because large size neurons bear a PNN in the DCN (Carulli et al., 2006; Hirono et al., 2018), which are excitatory (Uusisaari et al., 2007), only neurons with a size > 250 um^2^ (evaluated by ImageJ) were included in the analysis (uninjected, number of neurons = 14; injected, number of neurons = 16). GABAergic terminals were identified by the presence of oval/flattened synaptic vesicles and symmetric release sites. Glutamatergic terminals were identified by round synaptic vesicles and thick postsynaptic densities, which in some cases showed subsynaptic dense bodies, known as Taxi bodies (Uchizono, 1965; Kleim et al. 1998; Kleim et al. 2002; Fig. S4A, B). By means of ImageJ we evaluated: a) the size of individual GABAergic terminals around the cell body of IntA neurons (uninjected, number of terminals = 122; ch’ase, number of terminals = 145); b) the number of GABAergic terminals/mm soma membrane (uninjected, number of segments of neuronal membrane = 16; ch’ase, number of segments of neuronal membrane = 26); c) the distance between terminals (uninjected, number of pairs of terminals = 21; ch’ase, number of pairs of terminals = 41; d) the length of release sites (uninjected, number of release sites = 146; ch’ase, number of release sites = 152); e) the number of release sites/um of terminal membrane (uninjected, number of terminals = 72; ch’ase, number of terminals = 77); f) the area occupied by mitochondria/area of bouton (uninjected, number of terminals = 103; ch’ase, number of terminals = 90).

### In vivo *electrophysiology*

To investigate the effects of PNN removal on the electrophysiological properties of IntA neurons, mice were randomly assigned to a group receiving bilateral injections of LV-PGK-GFP (N = 6) or LV-PGK-ch’ase (N = 5) in the IntA nuclei (as described above). Three weeks after virus injection, mice were anesthetized using isoflurane and eyes were protected from drying using eye drops (Duodrops, Ceva). Body temperature was maintained at 37°C using a temperature-controlled heating pad guided by a rectal temperature probe. Skin in the neck was shaved and a 1 cm longitudinal incision was made to uncover the neck muscles. Muscles were locally anesthetized (10% Xylocaine, AstraZeneca) and removed to access the occipital bone. Two square craniotomies were made using a dental drill above Crus II, approximately 1.7 mm lateral to Bregma. A thin layer of primer (Optibond All-In-One, Kerr) was applied around the craniotomies, hardened out with UV light, and a circular recording bath was created using dental cement (Flowline and Charisma, Heraeus Kulzer). Bath was filled with saline and the dura was removed covering the brain in both craniotomies. Bath was then filled with a low viscosity silicone elastomer sealant (Kwik-cast, World Precision Instruments) to prevent the brain from drying and the skin surrounding the bath was attached to the skull bone with tissue glue (Histoacryl). Post-operative analgesia was given by i.p. injection of meloxicam (0.4 mg/kg Metacam) and mice recovered for at least 2 hours before initiation of the recording session. During recordings, mice were head-fixed. Extracellular recordings were performed using electrodes with a tip diameter of ~1 μm and a pipette resistance of 4-8 MΩ, heat-pulled from filamented borosilicate glass capillaries (1.5 mm OD, 0.86 mm ID, Harvard Apparatus) with a P-1000 micropipette puller (Sutter Instruments). The electrode was filled with standard intracellular solution, containing (in mM): 10 KOH, 3.48 MgCl_2_, 4 NaCl, 129 K-Gluconate, 10 hepes, 17.5 glucose, 4 Na_2_ATP, and 0.4 Na_3_GTP (295 ± 305 mOsm; pH 7.2). The intracellular solution was supplemented with 0.5% neurobiotin (Vector Labs). The electrode was attached to an electrode holder that approached the brain surface in a 43° angle and movements were controlled by a micromanipulator (SM7, Luigs and Neumann). Recordings were amplified using a Multiclamp700B amplifier (Axon Instruments) and digitized on 100 kHz using a Digidata 1440 (Axon Instruments). The electrode was lowered into the brain under high pressure till a depth of 1500 μm, after which the pressure was lowered to 15-20 mbar, and the electrode was advanced in steps of 2 μm per second until individual spikes could be identified with >1 mV amplitudes. The maximum duration of a recording session was 4 hours, after which mice were transcardially perfused for histology. Streptavidin-Cy3 (1:500, 1 h RT) and WFA FITC (Vector, 1:400, 1 h RT) were used to detect biocytin filled regions and PNNs, respectively.

### Electrophysiological data analysis

Spike times were obtained using a custom-written MATLAB spike analysis program (B.H.J. Winkelman, Netherlands Institute for Neuroscience, Amsterdam) and sorted based on spike waveform characteristics. In most recordings single units could be clearly identified. Recordings in which single units could not be unequivocally distinguished were excluded from analysis. Spontaneous spike characteristics were calculated based on 16.4 ± 7.8 seconds of recording while the mouse was in quiet wakefulness in the dark. As *in vivo* cell physiological parameters we analyzed: a) spike frequency (number of spikes/duration of time period); b) coefficient of variation of spiking [CV = standard deviation (all inter-spike intervals (ISIs)/average all ISIs)]; c) average CV of two adjacent ISIs [CV_2_ = 2 (ISI_n+1_ - ISI_n_)/(ISI_n_ + ISI_n_ + 1)].

### Statistical analysis

Statistical analysis was carried out by GraphPad Prism 5 and 7 (GraphPad Software Inc., La Jolla, CA, USA), SPSS version 22 (IBM) and MATLAB (R2011b, MathWorks). Normality of distributions was assessed using Shapiro-Wilk test and, if normality was violated, non-parametric tests were performed. Behavioural performance (EBC 5 days, EBC 10 days, and EBC retention phase) was analysed using repeated-measures ANOVA (data from camera setup) or linear mixed models (data from MDMT setup) with the maximum likelihood method. Group and session/day were modelled as fixed effects, and %CR and FEC at US onset as dependent variables. We assessed the fit of the model by running the analysis with the unstructured, diagonal and first-order autoregressive repeated covariance types, after which we choose the covariance type with the lowest Akaike’s information criterion value, which was in most cases the unstructured type. Chi-square test was used to compare the frequency distribution of WFA+ nets. Unpaired Student’s t-test or one-way ANOVA (followed by Tukey’s post-hoc test) were used to analyse synaptic terminals in immunohistochemistry- or electron microscopy-processed slices. Independent-samples Mann-Whitney U (MWU) test was used to compare spontaneous spike characteristics. Data are shown as mean ± SEM, and P ≤ 0.05 was considered as statistically significant.

## Supplemental information

**Figure S1. PNN reduction is further accentuated in the right DLH during EBC.** (A) Relative content of WFA+ nets in the DLH shows that the percentage of nets with strong WFA intensity is less in the right (R) DLH than in the left (L) DLH of conditioned (cond) mice on day 5 of training. The distribution of WFA+ nets in each side is different from that of CTR and pseudo mice. (B) In the lateral IntA no difference in WFA frequency distribution between right and left side is present in conditioned mice. The relative content of WFA+ nets in both the right (R) and the left (L) side of conditioned animals is different from that of CTR and pseudo animals. * P < 0.05, ** P < 0.01, *** P < 0.001.

**Figure S2. GABAergic and glutamatergic terminals in the DCN.** (A) Synaptic terminals in uninjected IntA, stained by antibodies against the general presynaptic marker SV2 (red) and VGAT (marker for GABAergic terminals; green). GABAergic terminals are the most abundant, and clearly outline neuronal somata (asterisks). In (B), the colocalisation (yellow) between calbindin (CALB, red), marker for Purkinje cells including their synaptic terminals, and VGAT (green) shows that the majority of terminals around DCN neurons belong to Purkinje cells. Arrows point to GABAergic terminals that do not belong to Purkinje cells (VGAT+/CALB-). CALB+ structures that do not express VGAT are Purkinje cell axons. Scale bar: 15 um in A, 5 um in B.

**Figure S3. Intensity profile of VGAT staining around DCN neurons.** (A-C) Representative plots showing intensity of VGAT fluorescence along the membrane of a neuronal soma in uninjected (A), LV-PGK-GFP injected (B) and LV-PGK-ch’ase injected IntA (C). The blue line indicates the arbitrary threshold selected for counting the number of troughs underneath it. On the “x” axis, the length of the neuronal membrane. (D) Quantification of the number of troughs/mm neuronal membrane. a.u. = arbitrary units. *** P < 0.001.

**Figure S4. EM pictures of GABAergic and glutamatergic terminals in the DCN.** (A) Example of GABAergic terminal, characterized by oval synaptic vesicles (black arrows) and symmetric release sites (open arrow). (B) Example of glutamatergic terminal, identified by round synaptic vesicles (black arrows). (B’) Glutamatergic terminal with thick post-synaptic density (open arrow). Scale bar: 200 nm in A (also applies to B); 400 nm in B’.

**Figure S5. Glutamatergic terminals are decreased in the DLH of LV-ch’ase injected mice during EBC retention.** (A, B) VGLUT1 immunolabeling in the DLH of LV-GFP (A) and LV-ch’ase (B) mice subjected to EBC and sacrificed after retention test. (C) Quantification of number of VGLUT1+ terminals/mm^2^. Scale bar: 50 um in A (also applies to B). * P < 0.05.

**Figure S6. Electrophysiological recording location.** (A) Insertion of neurobiotin-filled microstimulation electrode reveals neurobiotin labeling in the DCN. Arrows point to neurobiotin+ DCN neurons, as confirmed by MAP2 staining (B, C). The DCN is outlined by dashed lines (A-C). Scale bar: 100 um in A (also applies to B, C).

## References

Aizenman, C.D., and Linden, D.J. (1999). Regulation of the Rebound Depolarization and Spontaneous Firing Patterns of Deep Nuclear Neurons in Slices of Rat Cerebellum. J. Neurophysiol. 82, 1697–1709.

Bartus, K., James, N.D., Didangelos, A., Bosch, K.D., Verhaagen, J., Yáñez-Muñoz, R.J., Rogers, J.H., Schneider, B.L., Muir, E.M., and Bradbury, E.J. (2014). Large-Scale Chondroitin Sulfate Proteoglycan Digestion with Chondroitinase Gene Therapy Leads to Reduced Pathology and Modulates Macrophage Phenotype following Spinal Cord Contusion Injury. J. Neurosci. 34, 4822–4836.

Banerjee, S.B., Gutzeit, V.A., Baman, J., Aoued, H.S., Doshi, N.K., Liu, R.C., and Ressler, K.J. (2017). Perineuronal Nets in the Adult Sensory Cortex Are Necessary for Fear Learning. Neuron 95, 169–179.e3.

Bernard, C., and Prochiantz, A. (2016). Otx2-PNN Interaction to Regulate Cortical Plasticity. Neural Plast. 2016.

Beurdeley, M., Spatazza, J., Lee, H.H.C., Sugiyama, S., Bernard, C., Di Nardo, A.A., Hensch, T.K., and Prochiantz, A. (2012). Otx2 Binding to Perineuronal Nets Persistently Regulates Plasticity in the Mature Visual Cortex. J. Neurosci. 32, 9429–9437.

Blacktop, J.M., and Sorg, B.A. (2019). Perineuronal nets in the lateral hypothalamus area regulate cue-induced reinstatement of cocaine-seeking behavior. Neuropsychopharmacology 44, 850–858.

Blacktop, J.M., Todd, R.P., and Sorg, B.A. (2017). Role of perineuronal nets in the anterior dorsal lateral hypothalamic area in the acquisition of cocaine-induced conditioned place preference and self-administration. Neuropharmacology 118, 124–136.

Boele, H.-J., Koekkoek, S.K.E., De Zeeuw, C.I., and Ruigrok, T.J.H. (2013). Axonal Sprouting and Formation of Terminals in the Adult Cerebellum during Associative Motor Learning. J. Neurosci. 33, 17897–17907.

Boggio, E.M., Ehlert, E.M., Lupori, L., Moloney, E.B., De Winter, F., Vander Kooi, C.W., Baroncelli, L., Mecollari, V., Blits, B., Fawcett, J.W., et al. (2019). Inhibition of Semaphorin3A Promotes Ocular Dominance Plasticity in the Adult Rat Visual Cortex. Mol. Neurobiol.

Bosch, K.D., Bradbury, E.J., Verhaagen, J., Fawcett, J.W., and McMahon, S.B. (2012). Chondroitinase ABC promotes plasticity of spinal reflexes following peripheral nerve injury. Exp. Neurol. 238, 64–78.

Bowes, C., Massey, J.M., Burish, M., Cerkevich, C.M., and Kaas, J.H. (2012). Chondroitinase ABC promotes selective reactivation of somatosensory cortex in squirrel monkeys after a cervical dorsal column lesion. Proc. Natl. Acad. Sci. 109, 2595–2600.

Brodal, P., Dietrichs, E., and Walberg, F. (1986). Do pontocerebellar mossy fibres give off collaterals to the cerebellar nuclei? An experimental study in the cat with implantation of crystalline HRP-WGA. Neurosci. Res. 4, 12–24.

Brown, K.L., and Freeman, J.H. (2014). Extinction, reacquisition, and rapid forgetting of eyeblink conditioning in developing rats. Learn. Mem. 21, 696–708.

Brown, K.L., and Freeman, J.H. (2016). Retention of eyeblink conditioning in periweanling and adult rats. Dev. Psychobiol. 58, 1055–1065.

Burnside, E.R., De Winter, F., Didangelos, A., James, N.D., Andreica, E.C., Layard-Horsfall, H., Muir, E.M., Verhaagen, J., and Bradbury, E.J. (2018). Immune-evasive gene switch enables regulated delivery of chondroitinase after spinal cord injury. Brain 141, 2362–2381.

Carbo-Gas, M., Moreno-Rius, J., Guarque-Chabrera, J., Vazquez-Sanroman, D., Gil-Miravet, I., Carulli, D., Hoebeek, F., De Zeeuw, C., Sanchis-Segura, C., and Miquel, M. (2017). Cerebellar perineuronal nets in cocaine-induced pavlovian memory: Site matters. Neuropharmacology 125, 166–180.

Carmichael, S.T., Archibeque, I., Luke, L., Nolan, T., Momiy, J., and Li, S. (2005). Growth-associated gene expression after stroke: Evidence for a growth-promoting region in peri-infarct cortex. Exp. Neurol. 193, 291–311.

Carulli, D., Foscarin, S., Faralli, A., Pajaj, E., and Rossi, F. (2013). Modulation of semaphorin3A in perineuronal nets during structural plasticity in the adult cerebellum. Mol. Cell. Neurosci. 57, 10–22.

Carulli, D., Pizzorusso, T., Kwok, J.C.F., Putignano, E., Poli, A., Forostyak, S., Andrews, M.R., Deepa, S.S., Glant, T.T., and Fawcett, J.W. (2010). Animals lacking link protein have attenuated perineuronal nets and persistent plasticity. Brain 133, 2331–2347.

Carulli, D., Rhodes, K.E., Brown, D.J., Bonnert, T.P., Pollack, S.J., Oliver, K., Strata, P., and Fawcett, J.W. (2006). Composition of perineuronal nets in the adult rat cerebellum and the cellular origin of their components. J. Comp. Neurol. 494, 559–577.

Carulli, D., Rhodes, K.E., and Fawcett, J.W. (2007). Upregulation of aggrecan, link protein 1, and hyaluronan synthases during formation of perineuronal nets in the rat cerebellum. J. Comp. Neurol. 501, 83–94.

Chettih, S.N., McDougle, S.D., Ruffolo, L.I., and Medina, J.F. (2011). Adaptive Timing of Motor Output in the Mouse: The Role of Movement Oscillations in Eyelid Conditioning. Front. Integr. Neurosci. 5, 1–11.

Christian, K.M., and Thompson, R.F. (2003). Neural substrates of eyeblink conditioning: acquisition and retention. Learn. Mem. 10, 427–455.

Clark, R.E., Zhang, A.A., and Lavond, D.G. (1992). Reversible lesions of the cerebellar interpositus nucleus during acquisition and retention of a classically conditioned behavior. Behav. Neurosci. 106, 879–888.

Corvetti, L., and Rossi, F. (2005). Degradation of Chondroitin Sulfate Proteoglycans Induces Sprouting of Intact Purkinje Axons in the Cerebellum of the Adult Rat. J. Neurosci. 25, 7150–7158.

De Zeeuw, C.I., and Berrebi, A.S. (1995). Postsynaptic Targets of Purkinje Cell Terminals in the Cerebellar and Vestibular Nuclei of the Rat. Eur. J. Neurosci. 7, 2322–2333.

De Zeeuw, C.I., Hoebeek, F.E., Bosman, L.W.J., Schonewille, M., Witter, L., and Koekkoek, S.K. (2011). Spatiotemporal firing patterns in the cerebellum. Nat. Rev. Neurosci. 12, 327–344.

De Zeeuw, C.I., and Yeo, C.H. (2005). Time and tide in cerebellar memory formation. Curr. Opin. Neurobiol. 15, 667–674.

Déglon, N., Tseng, J.L., Bensadoun, J.C., Zurn, A.D., Arsenijevic, Y., Pereira de Almeida, L., Zufferey, R., Trono, D., and Aebischer, P. (2000). Self-inactivating lentiviral vectors with enhanced transgene expression as potential gene transfer system in Parkinson’s disease. Hum Gene Ther 11, 179–190.

Dick, G., Liktan, C., Alves, J.N., Ehlert, E.M.E., Miller, G.M., Hsieh-Wilson, L.C., Sugahara, K., Oosterhof, A., Van Kuppevelt, T.H., Verhaagen, J., et al. (2013). Semaphorin 3A binds to the perineuronal nets via chondroitin sulfate type E motifs in rodent brains. J. Biol. Chem. 288, 27384–27395.

Dull, T., Zufferey, R., Kelly, M., Mandel, R.J., Nguyen, M., Trono, D., and Naldini, L. (1998). A third-generation lentivirus vector with a conditional packaging system. J. Virol. 72, 8463–8471.

Duncan, J.A., Foster, R., and Kwok, J.C.F. (2019). The potential of memory enhancement through modulation of perineuronal nets. Br. J. Pharmacol.

Faralli, A., Dagna, F., Albera, A., Bekku, Y., Oohashi, T., Albera, R., Rossi, F., and Carulli, D. (2016). Modifications of perineuronal nets and remodelling of excitatory and inhibitory afferents during vestibular compensation in the adult mouse. Brain Struct. Funct. 221, 3193–3209.

Fawcett, J.W., Oohashi, T., and Pizzorusso, T. (2019). The roles of perineuronal nets and the perinodal extracellular matrix in neuronal function. Nat. Rev. Neurosci.

Foscarin, S., Ponchione, D., Pajaj, E., Leto, K., Gawlak, M., Wilczynski, G.M., Rossi, F., and Carulli, D. (2011). Experience-dependent plasticity and modulation of growth regulatory molecules at central synapses. PLoS One 6.

Freeman, J.H. Jr, Halverson, H.E., and Poremba, A. (2005). Differential effects of cerebellar inactivation on eyeblink conditioned excitation and inhibition. J. Neurosci. 25, 889–895.

Freeman, J.H. (2015). Cerebellar learning mechanisms. Brain Res. 1621, 260–269.

Freeman, J.H., and Steinmetz, A.B. (2011). Neural circuitry and plasticity mechanisms underlying delay eyeblink conditioning. Learn. Mem. 18, 666–677.

Frischknecht, R., Heine, M., Perrais, D., Seidenbecher, C.I., Choquet, D., and Gundelfinger, E.D. (2009). Brain extracellular matrix affects AMPA receptor lateral mobility and short-term synaptic plasticity. Nat. Neurosci. 12, 897–904.

Galtrey, C.M., Asher, R.A., Nothias, F., and Fawcett, J.W. (2007). Promoting plasticity in the spinal cord with chondroitinase improves functional recovery after peripheral nerve repair. Brain 130, 926–939.

Gao, Z., Van Beugen, B.J., and De Zeeuw, C.I. (2012). Distributed synergistic plasticity and cerebellar learning. Nat. Rev. Neurosci. 13, 619–635.

Gao, Z., Davis, C., Thomas, A.M., Economo, M.N., Abrego, A.M., Svoboda, K., Zeeuw, C.I. De, and Li, N. (2018). A cortico-cerebellar loop for motor planning. Nature.

Garcia, K.S., Steele, P.M., and Mauk, M.D. (1999). Cerebellar Cortex Lesions Prevent Acquisition of Conditioned Eyelid Responses. J. Neurosci. 19, 10940–10947.

Geissler, M., Gottschling, C., Aguado, A., Rauch, U., Wetzel, C.H., Hatt, H., and Faissner, A. (2013). Primary Hippocampal Neurons, Which Lack Four Crucial Extracellular Matrix Molecules, Display Abnormalities of Synaptic Structure and Function and Severe Deficits in Perineuronal Net Formation. J. Neurosci. 33, 7742–7755.

Gherardini, L., Gennaro, M., and Pizzorusso, T. (2015). Perilesional treatment with chondroitinase ABC and motor training promote functional recovery after stroke in rats. Cereb. Cortex 25, 202–212.

Gogolla, N., Caroni, P., Lüthi, A., and Herry, C. (2009). Perineuronal nets protect fear memories from erasure. Science (80-.). 325, 1258–1261.

Halverson, H.E., Khilkevich, A., and Mauk, M.D. (2015). Relating Cerebellar Purkinje Cell Activity to the Timing and Amplitude of Conditioned Eyelid Responses. J. Neurosci. 35, 7813–7832.

Happel, M.F.K., Niekisch, H., Castiblanco Rivera, L.L., Ohl, F.W., Deliano, M., and Frischknecht, R. (2014). Enhanced cognitive flexibility in reversal learning induced by removal of the extracellular matrix in auditory cortex. Proc. Natl. Acad. Sci. 111, 2800–2805.

Härtig, W., Brauer, K., and Brückner, G. (1992). Wisteria floribunda agglutinin-labelled nets surround parvalbumin-containing neurons. Neuroreport 3, 869–872.

Härtig, W., Derouiche, A., Welt, K., Brauer, K., Grosche, J., Mäder, M., Reichenbach, A., and Brückner, G. (1999). Cortical neurons immunoreactive for the potassium channel Kv3.1b subunit are predominantly surrounded by perineuronal nets presumed as a buffering system for cations. Brain Res. 842, 15–29.

Heiney, S.A., Wohl, M.P., Chettih, S.N., Ruffolo, L.I., and Medina, J.F. (2014). Cerebellar-Dependent Expression of Motor Learning during Eyeblink Conditioning in Head-Fixed Mice. J. Neurosci. 34, 14845–14853.

Hioki, H., Fujiyama, F., Taki, K., Tomioka, R., Furuta, T., Tamamaki, N., and Kaneko, T. (2003). Differential distribution of vesicular glutamate transporters in the rat cerebellar cortex. Neuroscience 117, 1–6.

Hirono, M., Watanabe, S., Karube, F., Fujiyama, F., Kawahara, S., Nagao, S., Yanagawa, Y., and Misonou, H. (2018). Perineuronal Nets in the Deep Cerebellar Nuclei Regulate GABAergic Transmission and Delay Eyeblink Conditioning. J. Neurosci. 38, 6130–6144.

Hou, X., Yoshioka, N., Tsukano, H., Sakai, A., Miyata, S., Watanabe, Y., Yanagawa, Y., Sakimura, K., Takeuchi, K., Kitagawa, H., et al. (2017). Chondroitin Sulfate Is Required for Onset and Offset of Critical Period Plasticity in Visual Cortex. Sci. Rep. 7, 1–17.

Hylin, M.J., Orsi, S.A., Moore, A.N., and Dash, P.K. (2013). Disruption of the perineuronal net in the hippocampus or medial prefrontal cortex impairs fear conditioning. Learn. Mem. 20, 267–273.

Ito M. (1972) Neural design of the cerebellar motor control system. Brain Res, 81–4.

Ivarsson, M., and Hesslow, G. (1993). Bilateral control of the orbicularis oculi muscle by one cerebellar hemisphere in the ferret. Neuroreport 4, 1127–1130.

Kandel, E.R., Dudai, Y., and Mayford, M.R. (2014). The molecular and systems biology of memory. Cell 157, 163–186.

Kistler, W.M., and Van Hemmen, J.L. (1999). Delayed reverberation through time windows as a key to cerebellar function. Biol. Cybern. 81, 373–380.

Kistler, W.M., van Hemmen, J.L., and De Zeeuw, C.I. (2000). Time window control: a model for cerebellar function based on synchronization, reverberation, and time slicing. Prog. Brain Res 124, 275–297.

Kleim, J.A., Freeman, J.H., Bruneau, R., Nolan, B.C., Cooper, N.R., Zook, A., and Walters, D. (2002). Synapse formation is associated with memory storage in the cerebellum. Proc. Natl. Acad. Sci. 99, 13228–13231.

Kleim, J.A., Pipitone, M.A., Czerlanis, C., and Greenough, W.T. (1998). Structural stability within the lateral cerebellar nucleus of the rat following complex motor learning. Neurobiol. Learn. Mem. 69, 290–306.

Knowlton, B.J., and Thompson, R.F. (1988). Microinjections of local anesthetic into the pontine nuclei reduce the amplitude of the classically conditioned eyelid response. Physiol. Behav. 43, 855–357.

Koekkoek, S.K.E., Den Ouden, W.L., Perry, G., Highstein, S.M., and De Zeeuw, C.I. (2002). Monitoring Kinetic and Frequency-Domain Properties of Eyelid Responses in Mice With Magnetic Distance Measurement Technique. J. Neurophysiol. 88, 2124–2133.

Krupa, D.J., and Thompson, R.F. (1997). Reversible inactivation of the cerebellar interpositus nucleus completely prevents acquisition of the classically conditioned eye-blink response. Learn. Mem. 3, 545–556.

Kwok, J.C., Afshari, F., García-Alías, G., and Fawcett, J.W. (2008). Proteoglycans in the central nervous system: plasticity, regeneration and their stimulation with chondroitinase ABC. Restor. Neurol. Neurosci. 26, 131–145.

Lee, H.H.C., Bernard, C., Ye, Z., Acampora, D., Simeone, A., Prochiantz, A., Di Nardo, A.A., and Hensch, T.K. (2017). Genetic Otx2 mis-localization delays critical period plasticity across brain regions. Mol. Psychiatry 22, 680–688.

Levy, C., Brooks, J.M., Chen, J., Su, J., and Fox, M.A. (2015). Cell-specific and developmental expression of lectican-cleaving proteases in mouse hippocampus and neocortex. J. Comp. Neurol. 523, 629–648.

Luque, N.R., Garrido, J.A., Carrillo, R.R., D’Angelo, E., and Ros, E. (2014). Fast convergence of learning requires plasticity between inferior olive and deep cerebellar nuclei in a manipulation task: a closed-loop robotic simulation. Front. Comput. Neurosci. 8, 1–16.

Mao, H., Hamodeh, S., and Sultan, F. (2018). Quantitative Comparison Of Vesicular Glutamate Transporters in rat Deep Cerebellar Nuclei. Neuroscience 376, 152–161.

Massey, J.M., Hubscher, C.H., Wagoner, M.R., Decker, J.A., Amps, J., Silver, J., Onifer, S.M. (2006). Chondroitinase ABC Digestion of the Perineuronal Net Promotes Functional Collateral Sprouting in the Cuneate Nucleus after Cervical Spinal Cord Injury. J. Neurosci. 26, 4406–4414.

McCornick, D.A., and Thompson, R.F. (1984). Neuronal responses of the rabbit cerebellum during acquisition and performance of a classically conditioned nictitating membrane-eyelid response. J. Neurosci. 4, 2811–2822

Miyata, S., Komatsu, Y., Yoshimura, Y., Taya, C., and Kitagawa, H. (2012). Persistent cortical plasticity by upregulation of chondroitin 6-sulfation. Nat. Neurosci. 15, 414–422.

Muir, E.M., Fyfe, I., Gardiner, S., Li, L., Warren, P., Fawcett, J.W., Keynes, R.J., and Rogers, J.H. (2010). Modification of N-glycosylation sites allows secretion of bacterial chondroitinase ABC from mammalian cells. J. Biotechnol. 145, 103–110.

Nakamura, H., Fujii, Y., Inoki, I., Sugimoto, K., Tanzawa, K., Matsuki, H., Miura, R., Yamaguchi, Y., and Okada, Y. (2000). Brevican is degraded by matrix metalloproteinases and aggrecanase-1 (ADAMTS4) at different sites. J. Biol. Chem. 275, 38885–38890.

Naldini, L., Blömer, U., Gallay, P., Ory, D., Mulligan, R., Gage, F.H., Verma, I.M., and Trono, D. (1996). In vivo gene delivery and stable transduction of nondividing cells by a lentiviral vector. Science 272, 263–267.

Nicholson, D.A., Sweet, J.A., and Freeman, J.H. (2003). Long-term retention of the classically conditioned eyeblink response in rats. Behav. Neurosci. 117, 871–875.

Paxinos, G., and Franklin, K.B.J., (2001). The mouse brain in stereotaxic coordinates. Academic press.

Perrett, S., Ruiz, B., and Mauk, M. (1993). Cerebellar cortex lesions disrupt learning-dependent timing of conditioned eyelid responses. J. Neurosci. 13, 1708–1718.

Pizzorusso, T., Medini, P., Berardi, N., Chierzi, S., Fawcett, J.W., and Maffei, L. (2002). Reactivation of ocular dominance plasticity in the adult visual cortex. Science (80-.). 298, 1248–1251.

Pugh, J.R., and Raman, I.M. (2009). Nothing can be coincidence: synaptic inhibition and plasticity in the cerebellar nuclei.

Rasmussen, A., Jirenhed, D. and Hesslow, G. (2008). Simple and Complex Spike Firing Patterns in Purkinje Cells During Classical Conditioning. 563–566.

Rasmussen, A., Ijpelaar, A.C.H.G., De Zeeuw, C.I., and Boele, H.J. (2018). Caffeine has no effect on eyeblink conditioning in mice. Behav. Brain Res. 337, 252–255.

Rochefort, C., Lefort, J., and Rondi-Reig, L. (2013). The cerebellum: a new key structure in the navigation system. Front. Neural Circuits 7, 1–12.

Romberg, C., Yang, S., Melani, R., Andrews, M.R., Horner, A.E., Spillantini, M.G., Bussey, T.J., Fawcett, J.W., Pizzorusso, T., and Saksida, L.M. (2013). Depletion of Perineuronal Nets Enhances Recognition Memory and Long-Term Depression in the Perirhinal Cortex. J. Neurosci. 33, 7057–7065.

Rowlands, D., Lensjø, K.K., Dinh, T., Yang, S., Andrews, M.R., Hafting, T., Fyhn, M., Fawcett, J.W., and Dick, G. (2018). Aggrecan Directs Extracellular Matrix-Mediated Neuronal Plasticity. J. Neurosci. 38, 10102–10113.

Sacchetti, B., Baldi, E., Lorenzini, C.A., and Bucherelli, C. (2002). Cerebellar role in fear-conditioning consolidation. Proc. Natl. Acad. Sci. U. S. A. 99, 8406–8411.

Sale, A., Maya Vetencourt, J.F., Medini, P., Cenni, M.C., Baroncelli, L., De Pasquale, R., and Maffei, L. (2007). Environmental enrichment in adulthood promotes amblyopia recovery through a reduction of intracortical inhibition. Nat. Neurosci. 10, 679–681.

Senkov, O., Andjus, P., Radenovic, L., Soriano, E., and Dityatev, A. (2014). Neural ECM molecules in synaptic plasticity, learning, and memory. Prog. Brain Res. 214, 53–80.

Slaker, M.L., Jorgensen, E.T., Hegarty, D.M., Liu, X., Kong, Y., Zhang, F., Linhardt, R.J., Brown, T.E., Aicher, S.A., and Sorg, B.A. (2018). Cocaine Exposure Modulates Perineuronal Nets and Synaptic Excitability of Fast-Spiking Interneurons in the Medial Prefrontal Cortex. Eneuro 5, ENEURO.0221-18.2018.

Slaker, M., Churchill, L., Todd, R.P., Blacktop, J.M., Zuloaga, D.G., Raber, J., Darling, R.A., Brown, T.E., and Sorg, B.A. (2015). Removal of Perineuronal Nets in the Medial Prefrontal Cortex Impairs the Acquisition and Reconsolidation of a Cocaine-Induced Conditioned Place Preference Memory. J. Neurosci. 35, 4190–4202.

Slaker, M., Blacktop, J.M., and Sorg, B.A. (2016). Caught in the net: Perineuronal nets and addiction. Neural Plast. 2016.

Smith, C.C., Mauricio, R., Nobre, L., Marsh, B., Wüst, R.C.I., Rossiter, H.B., and Ichiyama, R.M. (2015). Differential regulation of perineuronal nets in the brain and spinal cord with exercise training. Brain Res. Bull. 111, 20–26.

Soleman, S., Yip, P.K., Duricki, D.A., and Moon, L.D.F. (2012). Delayed treatment with chondroitinase ABC promotes sensorimotor recovery and plasticity after stroke in aged rats. 1210–1223.

Spatazza, J., Lee, H.H.C., DiNardo, A.A., Tibaldi, L., Joliot, A., Hensch, T.K., and Prochiantz, A. (2013). Choroid-Plexus-Derived Otx2 Homeoprotein Constrains Adult Cortical Plasticity. Cell Rep. 3, 1815–1823.

Stamenkovic, V., Stamenkovic, S., Jaworski, T., Gawlak, M., Jovanovic, M., Jakovcevski, I., Wilczynski, G.M., Kaczmarek, L., Schachner, M., Radenovic, L., et al. (2017). The extracellular matrix glycoprotein tenascin-C and matrix metalloproteinases modify cerebellar structural plasticity by exposure to an enriched environment. Brain Struct. Funct. 222, 393–415.

Takesian, A.E. and Hensch, T.K. (2013). Balancing plasticity/stability across brain development. Prog. Brain Res. 207, 3–34.

ten Brinke, M.M., Boele, H.J., Spanke, J.K., Potters, J.W., Kornysheva, K., Wulff, P., Ijpelaar, A.C.H.G., Koekkoek, S.K.E., and De Zeeuw, C.I. (2015). Evolving Models of Pavlovian Conditioning: Cerebellar Cortical Dynamics in Awake Behaving Mice. Cell Rep. 13, 1977–1988.

ten Brinke, M.M., Heiney, S.A., Wang, X., Proietti-Onori, M., Boele, H.J., Bakermans, J., Medina, J.F., Gao, Z., and De Zeeuw, C.I. (2017). Dynamic modulation of activity in cerebellar nuclei neurons during pavlovian eyeblink conditioning in mice. Elife 6, 1–27.

Thompson, E.H., Lensjø, K.K., Wigestrand, M.B., Malthe-Sørenssen, A., Hafting, T., and Fyhn, M. (2018). Removal of perineuronal nets disrupts recall of a remote fear memory. Proc. Natl. Acad. Sci. 115, 607–612.

Timmann, D., Drepper, J., Frings, M., Maschke, M., Richter, S., Gerwig, M., and Kolb, F.P. (2010). The human cerebellum contributes to motor, emotional and cognitive associative learning. A review. Cortex 46, 845–857.

Tortorella, M.D., Burn, T.C., Pratta, M.A., Abbaszade, I., Hollis, J.M., Liu, R., Rosenfeld, S.A., Copeland, R.A., Decicco, C.P., Wynn, R., et al. (1999). Purification and cloning of aggrecanase-1: a member of the ADAMTS family of proteins. Science 284, 1664–1666.

Tyagarajan, S.K., and Fritschy, J.M. (2014). Gephyrin: A master regulator of neuronal function? Nat. Rev. Neurosci. 15, 141–156.

Uchizono, K. (1965). Characteristics of excitatory and inhibitory synapses in the central nervous system of the cat. Nature 207, 642–643

Uusisaari, M., Obata, K., and Knöpfel, T. (2007). Morphological and Electrophysiological Properties of GABAergic and Non-GABAergic Cells in the Deep Cerebellar Nuclei. J. Neurophysiol. 97, 901–911.

Van Den Oever, M.C., Lubbers, B.R., Goriounova, N.A., Li, K.W., Van Der Schors, R.C., Loos, M., Riga, D., Wiskerke, J., Binnekade, R., Stegeman, M., et al. (2010). Extracellular matrix plasticity and GABAergic inhibition of prefrontal cortex pyramidal cells facilitates relapse to heroin seeking. Neuropsychopharmacology 35, 2120–2133.

Vazquez-Sanroman, D., Leto, K., Cerezo-Garcia, M., Carbo-Gas, M., Sanchis-Segura, C., Carulli, D., Rossi, F., and Miquel, M. (2015). The cerebellum on cocaine: Plasticity and metaplasticity. Addict. Biol. 20, 941–955.

Vo, T., Carulli, D., Ehlert, E.M.E., Kwok, J.C.F., Dick, G., Mecollari, V., Moloney, E.B., Neufeld, G., de Winter, F., Fawcett, J.W., et al. (2013). The chemorepulsive axon guidance protein semaphorin3A is a constituent of perineuronal nets in the adult rodent brain. Mol. Cell. Neurosci. 56, 186–200.

Vos, M., Lauwers, E., and Verstreken, P. (2010). Synaptic mitochondria in synaptic transmission and organization of vesicle pools in health and disease. Front. Synaptic Neurosci. 2, 1–10.

Wang, S.S., Kloth, A.D., and Badura, A. (2014). The cerebellum, sensitive periods, and autism. Neuron 83, 518–532.

Wetmore, D.Z., Mukamel, E.A., and Schnitzer, M.J. (2008). Lock-and-Key Mechanisms of Cerebellar Memory Recall Based on Rebound Currents. J. Neurophysiol. 100, 2328–2347.

Xue, Y.-X., Xue, L.-F., Liu, J.-F., He, J., Deng, J.-H., Sun, S.-C., Han, H.-B., Luo, Y.-X., Xu, L.-Z., Wu, P., et al. (2014). Depletion of Perineuronal Nets in the Amygdala to Enhance the Erasure of Drug Memories. J. Neurosci. 34, 6647–6658.

Yang, S., Cacquevel, M., Saksida, L.M., Bussey, T.J., Schneider, B.L., Aebischer, P., Melani, R., Pizzorusso, T., Fawcett, J.W., and Spillantini, M.G. (2015). Perineuronal net digestion with chondroitinase restores memory in mice with tau pathology. Exp. Neurol. 265, 48–58.

Yeo, C.H., Hardiman, M.J., and Glickstein, M. (1985). Classical conditioning of the nictitating membrane response of the rabbit. I. Lesions of the cerebellar nuclei. Exp. Brain Res. 60, 87–98.

Zheng, N., and Raman, I.M. (2009). Ca Currents Activated by Spontaneous Firing and Synaptic Disinhibition in Neurons of the Cerebellar Nuclei. J. Neurosci. 29, 9826–9838.

Zhao, R.R., Muir, E.M., Alves, J.N., Rickman, H., Allan, A.Y., Kwok, J.C., Roet, K.C.D., Verhaagen, J., Schneider, B.L., Bensadoun, J.C., et al. (2011). Lentiviral vectors express chondroitinase ABC in cortical projections and promote sprouting of injured corticospinal axons. J. Neurosci. Methods 201, 228–238.

