## Supplementary figures and images for "Cerebellar plasticity and associative memories are controlled by perineuronal nets"

### suppl fig 1

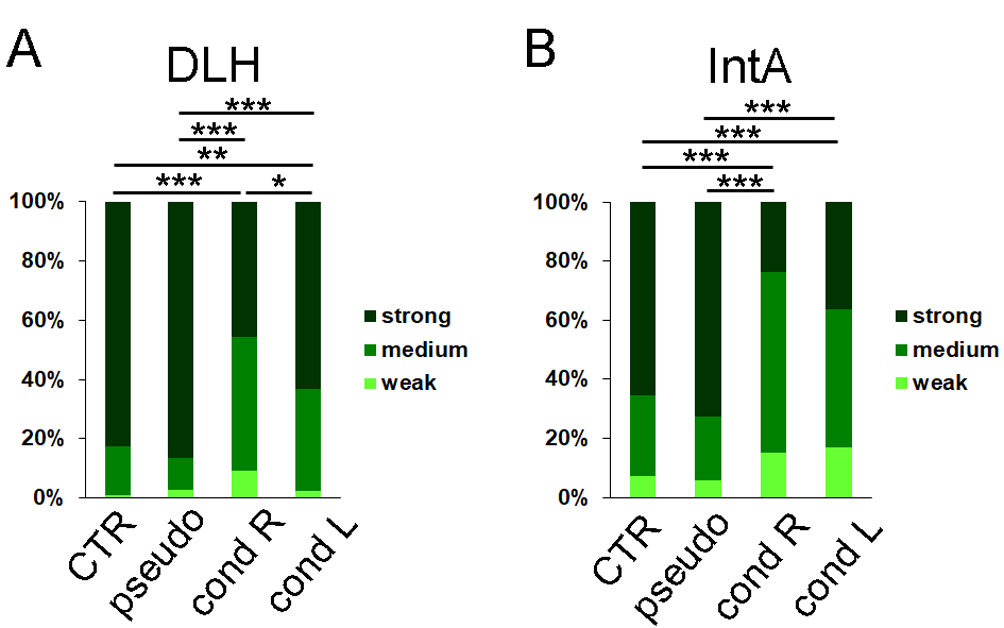

### suppl fig 2

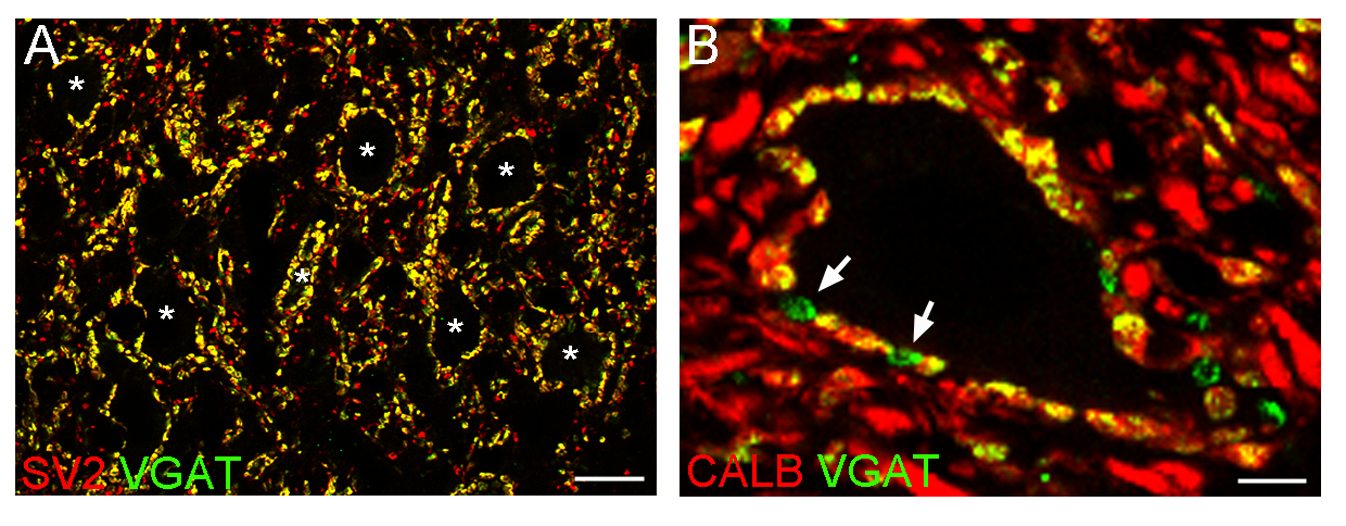

### suppl fig 3

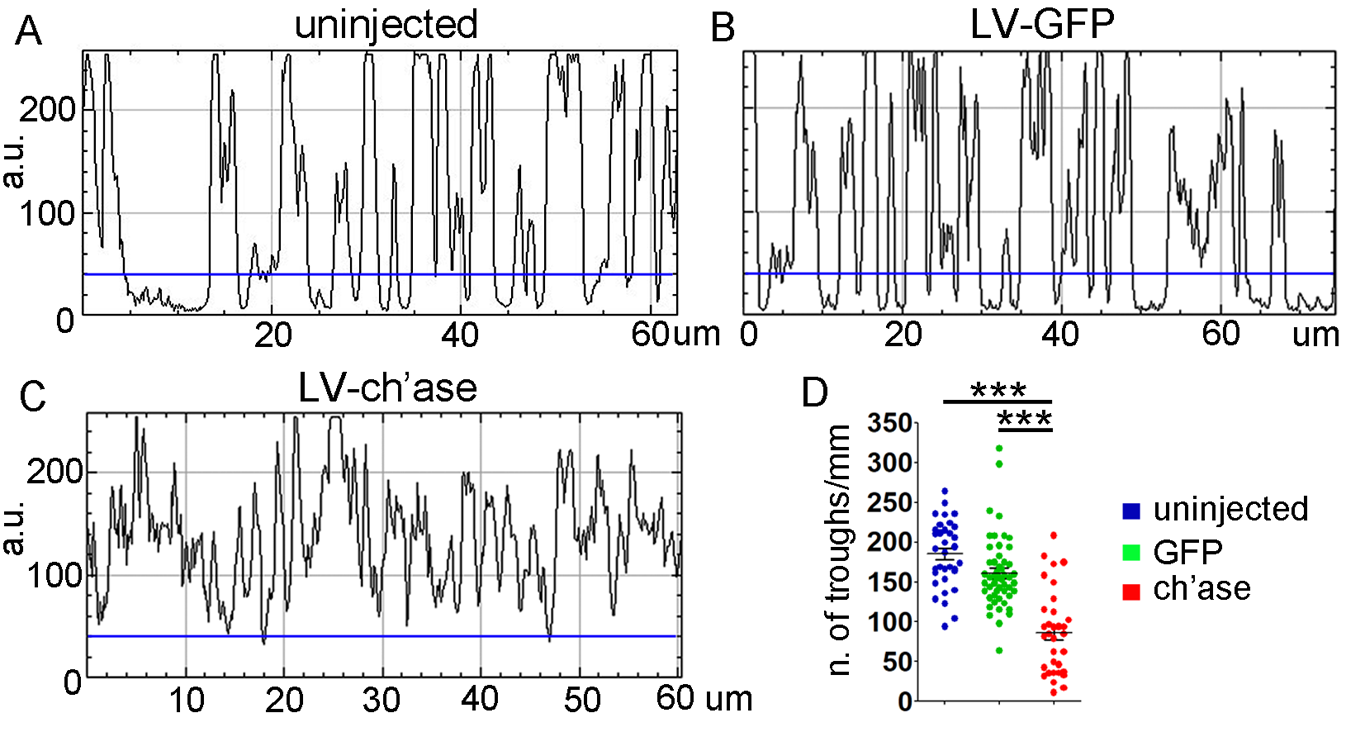

### suppl fig 4

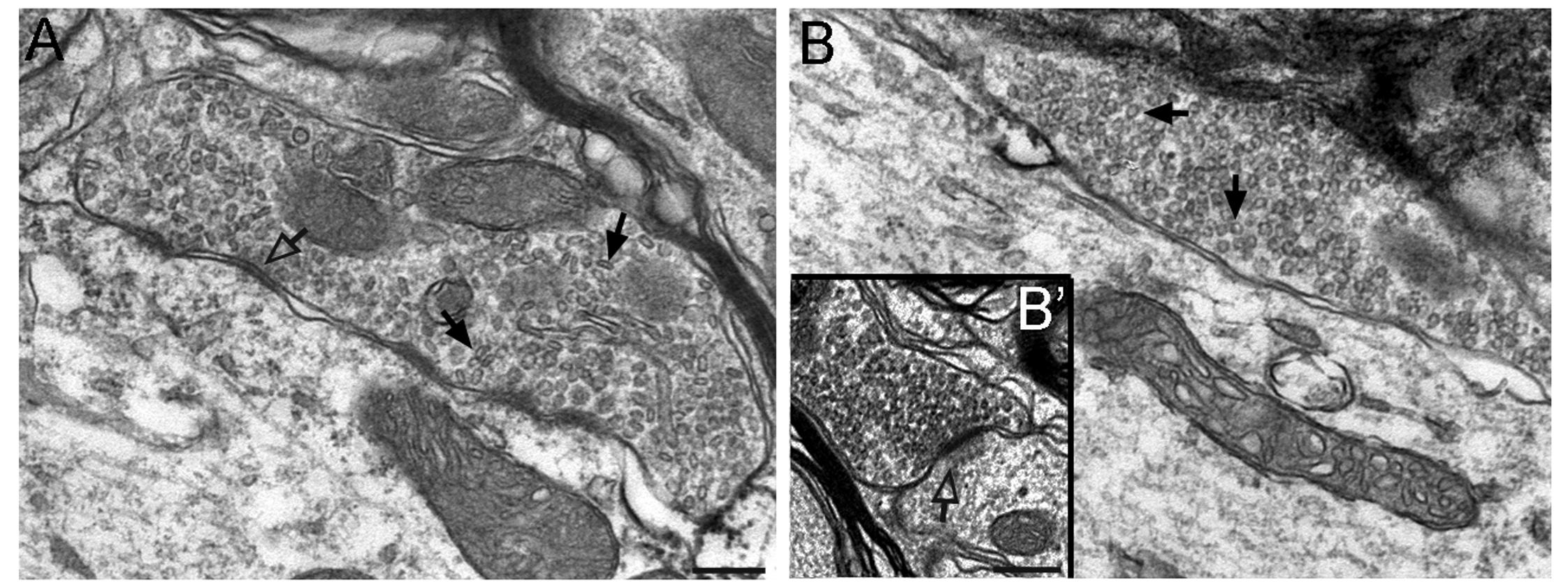

### suppl fig 5

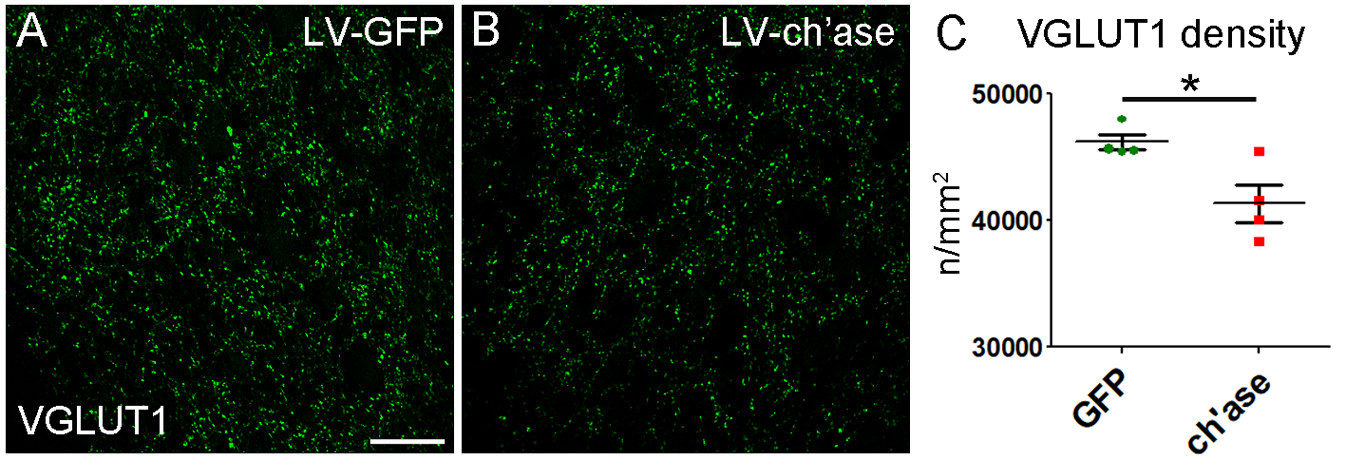

### suppl fig 6

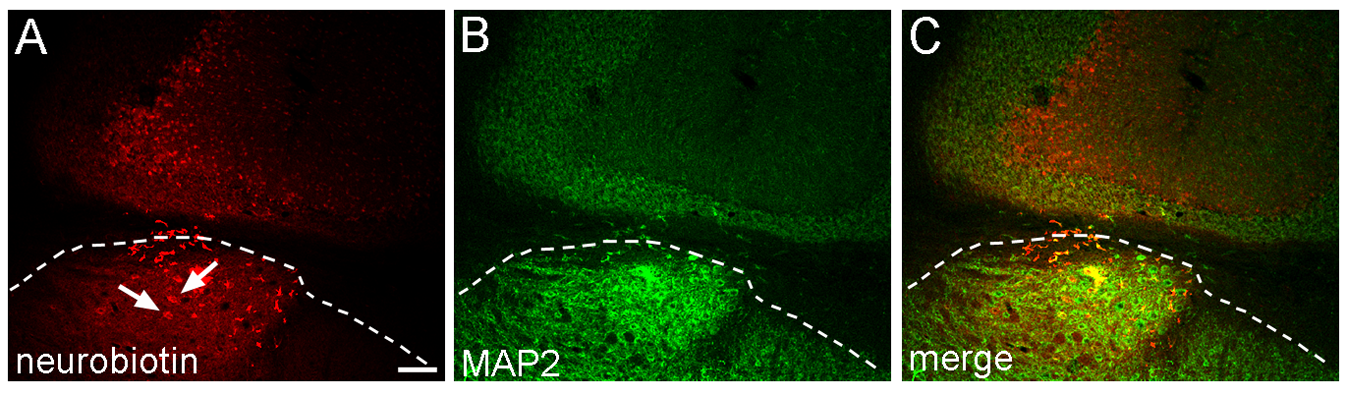
